# A human organoid model of alveolar regeneration reveals distinct epithelial responses to interferon-gamma

**DOI:** 10.1101/2025.01.30.635624

**Authors:** Antonella F.M. Dost, Katarína Balážová, Carla Pou Casellas, Lisanne M. van Rooijen, Wisse Epskamp, Gijs J.F. van Son, Willine J. van de Wetering, Carmen Lopez-Iglesias, Harry Begthel, Peter J. Peters, Niels Smakman, Johan. H. van Es, Hans Clevers

**Affiliations:** Hubrecht Institute - Royal Netherlands Academy of Arts and Sciences and University Medical Center Utrecht, Utrecht, The Netherlands; Oncode Institute, Utrecht, The Netherlands; Department of Nephrology and Hypertension, University Medical Center Utrecht, Utrecht, The Netherlands; The Princess Máxima Center for Pediatric Oncology, Utrecht, The Netherlands; Microscopy CORE Lab, Faculty of Health, Medicine and Life Sciences, Maastricht University, Maastricht, The Netherlands; The Maastricht Multimodal Molecular Imaging Institute, Maastricht University, Maastricht, The Netherlands; Department of Surgery, Diakonessenhuis, Utrecht, The Netherlands

## Abstract

Chronic obstructive pulmonary disease is characterized by inflammation and emphysema, leading to progressive alveolar destruction. Currently, no therapies effectively regenerate the alveolar epithelium. Here, we describe a feeder- and serum-free primary adult human organoid model to investigate how inflammation influences alveolar regeneration. We achieve long-term expansion of multipotent progenitor-like cells, while an alveolar type 2 (AT2) maturation protocol enhances surfactant production and supports tubular myelin formation. Introducing a LATS inhibitor to the expansion condition induces alveolar type 1 (AT1) differentiation while maintaining AT2 cells. Using this platform, we find that interferon-gamma exerts cytotoxic effects on AT1 cells while promoting the growth of regenerating AT2 cells, illustrating how a single inflammatory stimulus can have divergent effects on alveolar epithelial cell types. These findings underscore the nuanced influence of pro-inflammatory cytokines on alveolar regeneration. Our organoid model provides a reductionist platform for mechanistic studies, aimed to identify strategies to enhance alveolar regeneration.

## Introduction

Chronic obstructive pulmonary disease (COPD) is the third leading cause of death worldwide, with rising incidence^1^. The primary risk factor for developing COPD is chronic exposure to airborne toxic particles, such as cigarette smoke, air pollutants, dust, fumes, and chemicals^2^. The pathophysiology of COPD is complex and poorly understood, and involves airway obstruction, mucociliary dysfunction, chronic inflammation, and emphysema, the destruction of the alveolar region^3^. These features persist even after exposure cessation, making COPD a progressive disease with no cure. Available treatments, such as bronchodilators and corticosteroids, can alleviate shortness of breath and reduce inflammation. However, inhibiting the inflammatory response can increase the susceptibility to pulmonary infections, a serious complication that can lead to disease exacerbation. Moreover, to date there are no therapies that succeed in regenerating the damaged lung alveolar tissue and reversing emphysema^4^.

Lung alveoli are the gas-exchanging units of the lungs. Two major epithelial cell types exist within the alveoli: alveolar type 2 (AT2) and type 1 (AT1) cells. AT2 cells produce pulmonary surfactant that lines the alveolar epithelium. This surfactant, comprised of lipids and proteins, prevents collapsing of the alveoli by decreasing surface tension and plays a role in innate immunity. AT1 cells are exceedingly thin and spread out, forming part of the air-blood-barrier where gas exchange occurs. Not much is known about how human lungs regenerate, as our research relies heavily on mouse models. While AT2 cells are generally thought of as stem cells that can self-renew and differentiate to AT1 cells^5–7^, recent advances in single-cell RNA-Sequencing (scRNA-Seq) have helped define new cell types or cell states that might play a role in alveolar regeneration. Most noteworthy are the discovery of the AT2-signaling (AT2-s) population^8^, and cells residing at the terminal and respiratory bronchioles, termed pre-terminal bronchioles (pre-TB) secretory, TRB-specific alveolar type-0 (AT0)^9^, and respiratory airway secretory (RAS) cells^10^. One commonality of those populations is that they express lower levels of canonical AT2 markers and higher levels of secretory markers such as secretoglobin family 3A member 2 (SCGB3A2) as compared to the bulk AT2 population. This is in line with alveolar regeneration in murine models, where bronchioalveolar stem cells and club cells have been shown to be able to repair the alveolar epithelium^11–14^. Moreover, both in mouse models and in humans it has been shown that an AT2-AT1 transitional cell state emerges during injury repair^15–18^. Compared to AT2 cells, AT0 cells have higher expression of AT1 markers, placing them in between AT2, AT1, and secretory cells^9^. How these independently identified populations relate to each other is not completely understood. Even less understood are the roles of these populations and the mechanisms of alveolar regeneration in disease contexts such as COPD and emphysema, a knowledge gap that is hindering the development of new therapies. To address this unmet therapeutical need, it is important to develop human models that accurately recapitulate alveolar regeneration in the context of inflammatory disease.

Organoids are three-dimensional structures derived from stem cells grown in basement membrane-extract (BME), enabling us to study stem cell biology of untransformed human cells in vitro. Organoids are grown in conditions that promote proliferation and stemness. Because the lungs are quiescent during homeostasis, lung organoid cultures can be seen as an injury model that recapitulates regeneration rather than homeostasis. In recent years, primary feeder-free human adult alveolar organoid (ALVO) models have emerged, greatly advancing our toolbox to study alveolar regeneration^19–21^. However, these models were not characterized at single cell level and have a few drawbacks. Firstly, they rely on the surface marker HTII-280 to enrich for AT2 cells from whole lung tissue. However, it is unknown if a positive selection for HTII-280+ excludes cells that could give rise to alveolar cells in human lungs under regeneration conditions, such as the AT2-s, AT0, or RAS cells. Secondly, the protocols rely on serum to generate AT1 cells and hence do not provide defined media conditions, a crucial requirement for controlled disease modeling and mechanistic studies. In this study, we have aimed to develop a robust human ALVO model to overcome these challenges.

## Results

### Optimized ALVO conditions allow for long-term expansion in defined media

Current ALVO culturing protocols rely on positive selection of HTII-280+ AT2 cells (figure 1A). However, it is unknown whether this strategy excludes cells that are important for alveolar regeneration. To avoid excluding this population in our ALVO cultures and to maximize cell outgrowth, we deviated from existing protocols (figure 1A). We processed distal non-tumor human lung tissue from lobectomies of lung cancer patients into near single-cell suspensions and plated all cells in submerged, serum-free organoid conditions (SFFF media^19^; table S1). The omission of cell sorting shortens the processing time of the tissue and puts less stress on the cells. Furthermore, the presence of non-epithelial cells at passage (p)0 may support organoid outgrowth. After 14 days, large organoids with either alveolar (SFTPC) or airway (KRT5) markers had formed (figure 1B). Unlike organoids grown in airway organoid (AO) media^22^, the KRT5+ organoids appeared dense and did not express the club cell marker SCGB1A1^22^. We then enriched for epithelial cells and excluded airway progenitor cells by sorting the organoids for cells that expressed epithelial cell adhesion molecule (EPCAM) but not nerve growth factor receptor (NGFR) using fluorescence-activated cell sorting (FACS) (figure 1A and S1A). With some variation between organoid lines, the EPCAM+ fraction contained around 90% NGFR-cells (figure S1B). These ≥p1 EPCAM+/NGFR-cells gave rise to organoids that did not stain for airway markers (KRT5, SCGB1A1), while staining for an alveolar marker (SFTPC) (figure 1B). Notably, the cultures also contained cells and whole organoids that appeared to be negative for all three markers (figure 1B).

**Figure 1:**
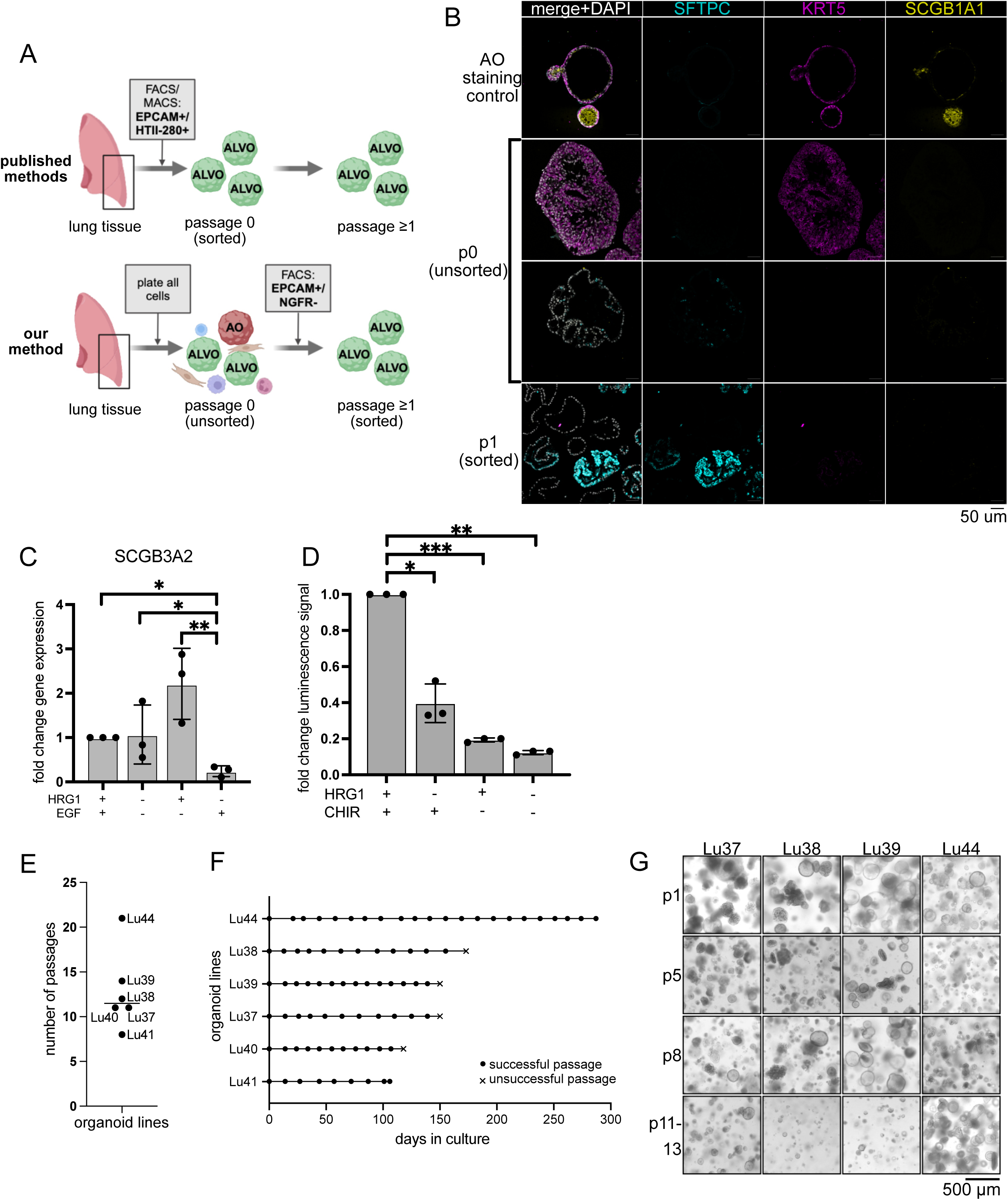
Optimized ALVO conditions allow for long-term expansion in defined media. A) Schematic comparing the published method and our method of generating ALVOs from lung tissue. B) IF images of AO organoids raised in AO media as a staining control, p0 organoids generated with our method, and p1 organoids after EPCAM+/NGFR-FACS sorting as described in 1A. DAPI = nuclei; SFTPC = AT2 markers; KRT5 = basal cell marker; SCGB1A1 = club cell marker. C) qPCR analysis of SCGB3A2 in ALVOs cultured in indicated media conditions for 14 days. Data are represented as mean ± SD. D) Viability assay (cell titer glo) of ALVOs cultured with indicated factors for 14 days. Data are represented as mean ± SD. E) Overview of number of passages reached for indicated ALVO lines. Median is indicated by a horizontal line. F) Overview of passage timings for indicated ALVO lines. G) Brightfield images of indicated ALVO lines at indicated passages. See also figure S1.

Next, we set out to improve long-term culturing conditions by refining the media composition. Specifically, we tested if we could replace epidermal growth factor (EGF) in the media with heregulin-beta-1 (HRG1). EGF and HRG1 bind to different members of the EGF receptor (EGFR) family, and EGF can lead to reduced long-term outgrowth in non-lung organoid systems^23^. Moreover, HRG1 has a positive effect on the growth and survival of airway organoids^24^. While we did not see a large difference between EGF and HRG1 in well-growing organoid cultures, we noticed that organoids at high passages (≥p7) appeared healthier with HRG1 compared to EGF (figure S1C). There was no difference in gene expression levels of the AT2 marker *SFTPC* and the AT1 marker *AGER* between media containing EGF or HRG1 after 14 days, indicating that EGF can be replaced with HRG1 without altering alveolar marker gene expression (figure S1D). Notably, the expression of *SCGB3A2* was higher in organoid growing in media containing HRG1 than in media containing EGF, possibly indicating a more progenitor-like state of the cells (figure 1C). Given this data, we replaced EGF with HRG1 and arrived at our alveolar organoid expansion media (ALVO-EM; table S1).

We tested ALVO-EM side-by-side with the commercially available alveolar organoid expansion (AvOE) media from Stem Cell Technologies (SCT). We found that after 14 days, the cells had a ∼30-fold reduced gene expression of *SFTPC* and an ∼11-fold higher expression of *CAV1* in AvOE, indicating increased AT1 differentiation compared to our ALVO-EM (figure S1E). We concluded that our media was better suited to prevent differentiation and promote expansion of AT2 cells within ALVOs.

Because the MAPK and Wnt pathways have been shown to be important drivers of proliferation in alveolar cells, we tested if this was also the case in our ALVOs. Withdrawal of either HRG1, CHIR99021 (CHIR; GSK3-beta inhibitor/Wnt activator), or both from the ALVO-EM led to a significant reduction in cell numbers, indicating that indeed both pathways are essential for ALVO growth (figure 1D).

Using the above-described strategy (figure 1A), we derived ALVO lines from multiple patients and passaged them in ALVO-EM conditions every 10-15 days. We achieved a median of 11 passages, with our best growing line still expanding at passage 21 (figure 1E-G). We monitored gene expression of alveolar markers over multiple passages of 4 of the lines (Lu37, Lu38, Lu39, Lu44) and found that expression levels of AT2 markers were stably high, mostly above expression levels of the house keeping gene beta-actin, while levels of AT1 marker *AGER* remained low in the first 10 passages (figure S1F). Therefore, we conducted all further experiments within the first 10 passages, mostly using Lu37, Lu38, and Lu39 as three biological replicates.

### Generating fully mature AT2 cells that secrete tubular myelin-containing surfactant

The ALVO-EM was developed to sustain long-term proliferation and self-renewal while still maintaining the AT2-like phenotype. However, we observed organoids with low or no protein expression of SFTPC, indicating that they might not contain fully mature AT2 cells (figure 1B). This was in line with the observation that the putative alveolar stem cell population in humans expresses lower levels of surfactant proteins^8–10^. To develop AT2-maturation media (AT2-MM) conditions, we screened for factors that increased the retention of lysotracker dye in the cells after initial organoid outgrowth (figure 2A). Lysotracker retention correlates with AT2 maturity, as it accumulates in the surfactant-producing and -storing organelles of the AT2 cell, the lamellar bodies^25^. We found that the addition of dexamethasone, cyclic adenosine monophosphate, and 3-isobutyl-1-methylxanthine, a cocktail termed “DCI”, used in induced pluripotent stem cell (iPSC)-derived organoids to induce AT2 maturation^26,27^, gave us the highest increase in lysotracker signal, followed by the cytokine interleukin (IL)-6 (figure 2B). The combined addition of DCI and IL-6 led to a higher increase in lysotracker signal compared to the single treatments (figure S2A and S2B). To further confirm the increased maturation state of the AT2 cells, we performed electron microscopy and observed not only microvilli and lamellar bodies, but also pulmonary surfactant secreted into the lumen of the organoids (figure 2C). Remarkably, the secreted surfactant formed highly ordered structures, including mesh-like tubular myelin, which has important innate immunity functions (figure 2C)^28^. However, when we analyzed expression levels of genes coding for proteins important for surfactant production (*LAMP3*, *NAPSA*), and surfactant proteins (*SFTPA1*, *SFTPD*, *SFTPC*), we found that - while the addition of DCI alone caused an increase or no change in all markers-, the addition of IL6 alone or in combination with DCI caused a decrease in *SFTPC* (figure S2C). Because *SFTPC* is important for the integrity of the surfactant lipid layer, we decided not to include IL6 in our AT2-MM (table S1) going forward. To confirm the increased AT2 maturity of the ALVOs in AT2-MM, we stained the surfactant-containing lamellar bodies of the organoids with the lipid dye Nile red, as pulmonary surfactant comprises of 90% lipids. As expected, we observed an increase in Nile red in AT2-MM compared to ALVO-EM (figure 2D and 2E). Notably, organoids with high Nile red signal did not necessarily stain for HTII-280, further emphasizing that the relationship between HTII-280 staining and AT2 status needs further investigation (figure 2D).

**Figure 2:**
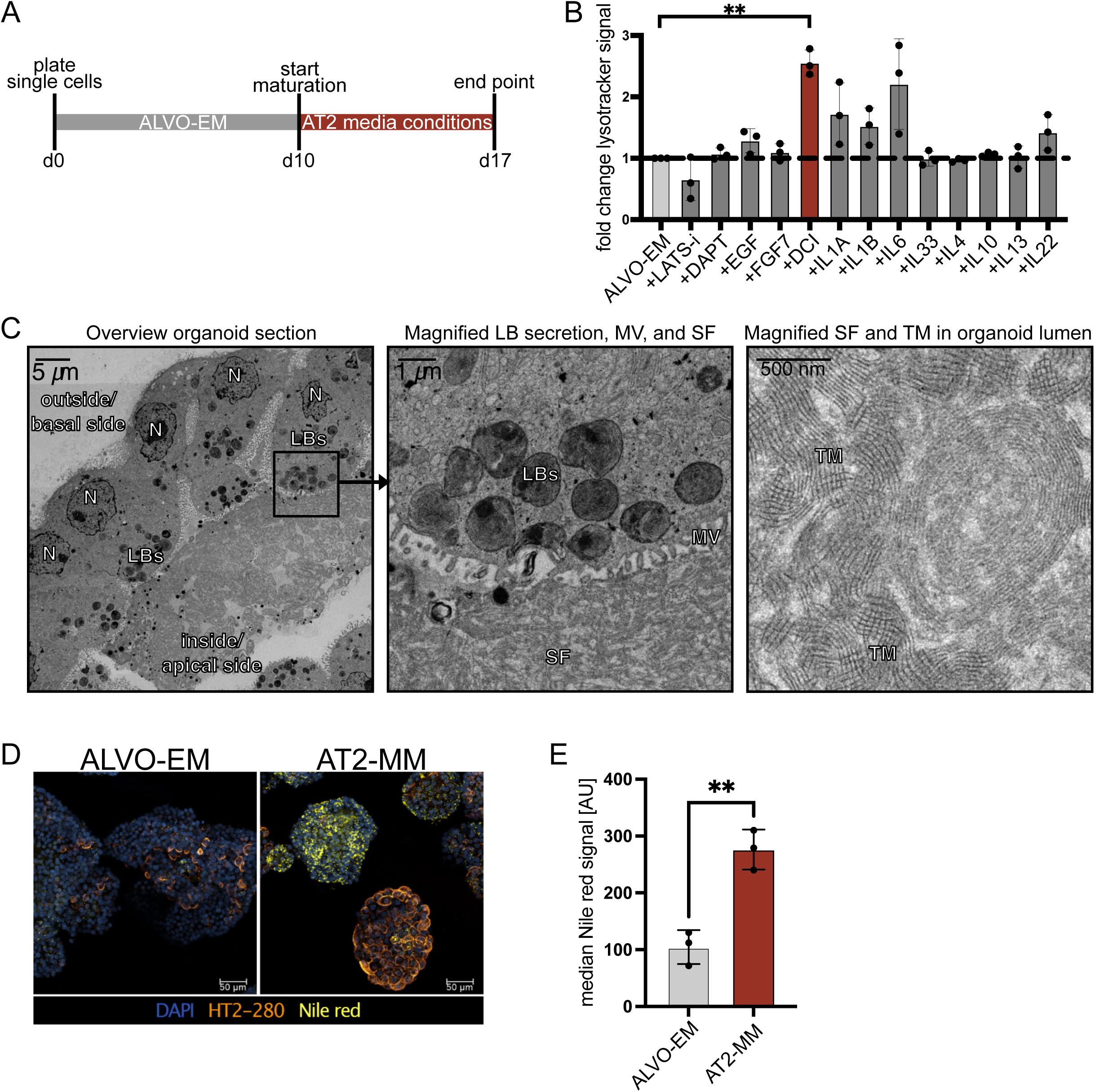
Generating fully mature AT2 cells that secrete tubular myelin-containing surfactant. A) Time line of media conditions used for ALVO cultures with indicated days (d). B) Flow cytometry analysis of median lysotracker signal of organoids treated with indicated factors as outlined by the timeline in 2A. Data are represented as mean ± SD. C) Electron microscopy image of ALVOs exposed to DIC and IL6 as outlined in 2A. Left: Part of an organoid containing mature AT2 cells and secreted surfactant inside the lumen. Center: Magnification of the apical side of one AT2 cell, showing MV and the process of the secretion of a LB into the lumen. Right: Magnification of the secreted SF and TM in the lumen of the organoid. N=nucleus; LBs=lamellar bodies; SF=surfactant; MV=microvilli; TM=tubular myelin. D) IF image of d17 organoids in ALVO-EM or AT2-MM as outlined in 2A. Nile red = lipid stain; HT2-280 = AT2 marker. E) Flow cytometry analysis showing median Nile red signal of d17 organoids in ALVO-EM or AT2-MM as outlined in 2A. Data are represented as mean ± SD. See also figure S2.

### Removal of proliferation factors and inhibition of LATS drives AT1 differentiation

To study alveolar regeneration in a controlled in vitro environment, defining media conditions to induce AT1 differentiation is crucial. Existing protocols use human serum (HS) to induce differentiation^29^. In iPSC-derived organoids, it was shown that a LATS-inhibitor (LATS-i) drives the AT1 phenotype^30^. To develop an AT1 differentiation medium, we removed all factors from the ALVO-EM that support proliferation and stemness to create a minimal base media (BM) (table S1). After expansion of the ALVOs for 10 days, we switched to the SCT differentiation media (AvOD), BM, BM + LATS-i, and BM + HS for 7 days (figure 3A). We found that the BM was sufficient to induce the upregulation of AT1 markers (*AGER, CAV1*) and downregulation of AT2 markers (*SFTPA1, SFTPC*) (figure S3A). This effect was strongest in the BM + LATS-i condition, indicating that the removal of proliferation and stemness factors in combination with LATS inhibition induced the AT1 program more strongly than HS and the commercial SCT media. Organoids in BM + LATS-i were smaller and more compact compared to ALVO-EM and AT2-MM (figure 3B). When we tested YAP target gene expression^31^, we found that *ANKRD1* and *CCN1* were upregulated in BM + LATS-i compared to ALVO-EM, suggesting that the differentiation was indeed correlated with active YAP signaling (figure S3B). Notably, BM alone was sufficient to increase YAP target genes, indicating that the default AT1 differentiation trajectory of AT2 cells is concomitant with active YAP signaling (figure S3B). To investigate the differentiation dynamics, we performed a qPCR time course for 14 days after switching to BM + LATS-i (figure S3C). Expression levels of the AT2 marker *SFTPC* already decreased after 2 days in BM + LATS-i and kept decreasing over time. The AT1 markers *AGER* and *CAV1* increased after 2 days of differentiation and kept increasing until they reached a plateau around 7 days. When we stained the organoids for SFTPC and AGER protein, we found that SFTPC was completely gone, while all organoids expressed high levels of AGER (figure S3D). Electron microscopy confirmed that the cells did not contain lamellar bodies (figure S3E). Instead, we observed cell shapes that appeared to have a low cytoplasm-to-nucleus ratio, reminiscent of AT1 cells. We did not observe the typical elongated AT1 cell shape in our 3D organoids. Instead, we observed large vacuoles in the cells, a sign of apoptosis and an unhealthy culture (figure S3E). Furthermore, these organoids were very difficult to dissociate enzymatically and mechanically without harming the cells, making analysis that requires single-cell suspensions challenging.

**Figure 3:**
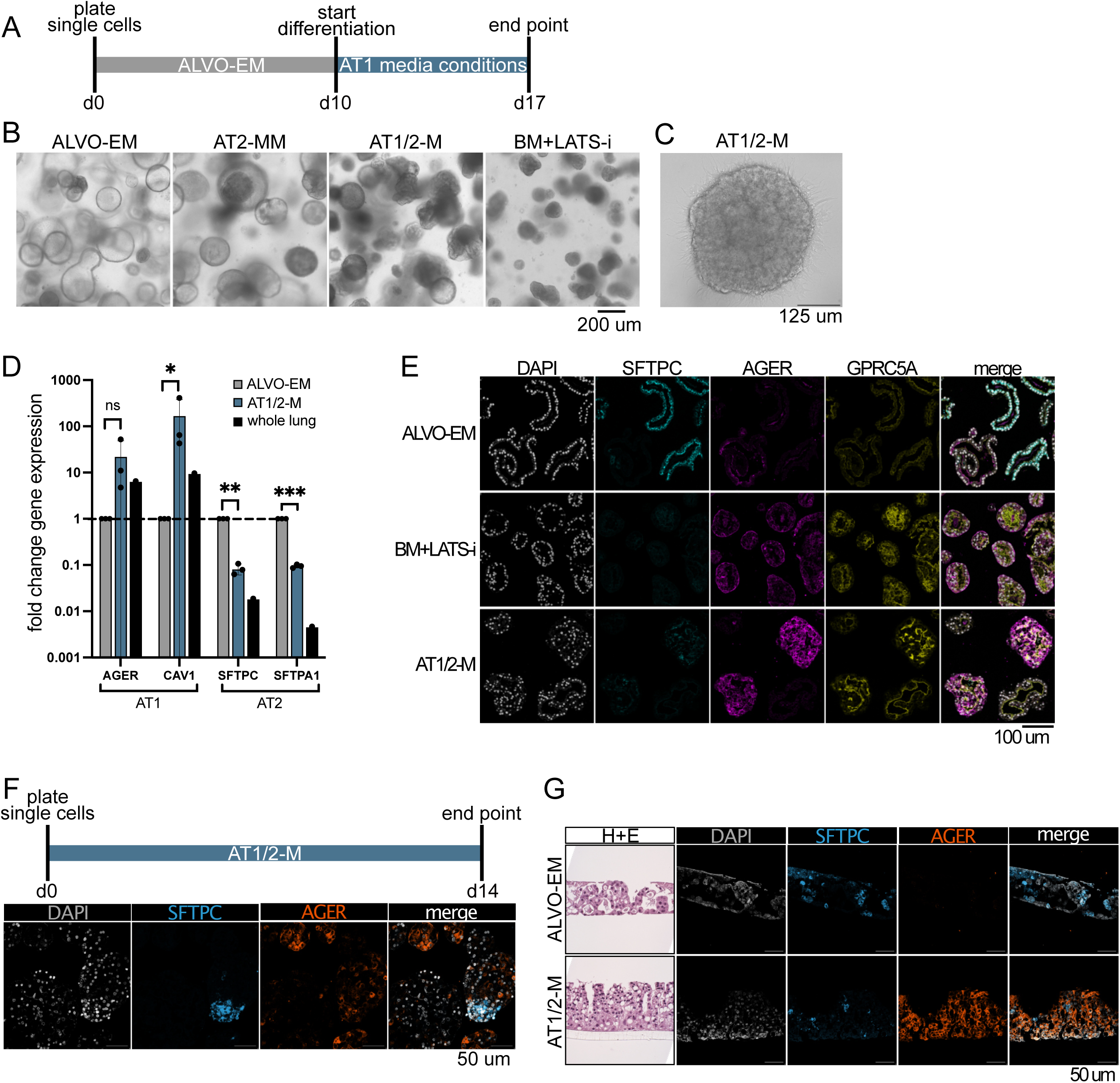
Addition of LATS-i to expansion media results in mixed AT1/AT2 organoids. A) Time line of media conditions used for ALVO cultures with indicated days (d). B) Brightfield images of p17 ALVO organoids cultured in indicated conditions as outlined in 3A. C) Close-up brightfield image of organoid cultured in AT1/2-M as outlined in 3A with visible attachment to the plate. D) qPCR analysis of AT1 (*AGER* and *CAV1*) and AT2 markers (*SFTPA1* and *SFTPC*) in ALVOs cultured in indicated conditions as outlined in 3A. Data are represented as mean ± SD. E) IF images of ALVOs cultured in indicated conditions as outlined in 3A. DAPI = nuclei; SFTPC = AT2 marker; AGER and GPRC5A = AT1 marker. F) IF images of ALVOs cultured in AT1/2-M for 14 days as indicated by time line. DAPI = nuclei; SFTPC = AT2 marker; AGER = AT1 marker. G) Side-view brightfield (H+E staining) and IF images of cells in 2D ALI transwells cultured in indicated media conditions. DAPI = nuclei; SFTPC = AT2 marker; AGER = AT1 marker. See also figure S3.

### Addition of LATS-i to expansion media results in mixed AT1/AT2 organoids

We concluded that conditions that allow for both AT1 and AT2 cells to be present at the same time might lead to more physiologically relevant cultures. Therefore, we tested whether adding LATS-i to our ALVO-EM (=AT1/2-M; table S1) is sufficient to drive AT1 differentiation in some cells. After an initial expansion phase of 10 days, we added LATS-i for 7 days and observed a striking change in morphology, with darker, more compact organoids (figure 3B) and increased attachment to the plate and growth in 2D (figure 3C). In contrast to BM + LATS-i, these organoids were not smaller than organoids in ALVO-EM. Gene expression analysis confirmed an increase in AT1 markers and a decrease in AT2 markers compared to ALVO-EM (figure 3D). IF staining showed that the organoids contained SFTPC-expressing AT2 cells, and cells with strong expression of the AT1 markers AGER and GPRC5A, indicating a mix of both cell types in the AT1/2-M culture conditions (figure 3E). Because we did not observe a decrease in size of the organoids grown in AT1/2-M compared to ALVO-EM, we tested whether the organoids would grow in AT1/2-M added from day 0 of culture, when they were still single cells (figure 3F). Surprisingly, the organoids grew out well and stained for SFTPC and AGER after 14 days (figure 3F), indicating that inhibition of LATS triggers AT1 differentiation without inhibiting proliferation.

In airway cells, two-dimensional (2D) air-liquid-interface (ALI) conditions induce differentiation^22^. Therefore, we tested if these conditions were sufficient to trigger AT1 differentiation in ALVOs. After 7 days of ALI culture with ALVO-EM at the bottom, we observed structured 3D growth. However, we did not detect AGER protein, indicating that 2D ALI cultures alone were not sufficient to induce AT1 marker expression (figure 3G). We tested if AT1/2-M would lead to AGER induction and found that—similar to our 3D cultures—we detected SFTPC+ and AGER+ cells in ALI cultures (figure 3G). Despite strong AGER expression, we did not observe the typical AT1 shape as found in vivo. Therefore, we grew the organoids in AT1/2-M conditions before we plated the cells in 2D ALI cultures (figure S3F). Because cell adhesion plays an important role in maintaining the AT1 cell shape, we tested different coatings of the transwells, including BME, invasin^32^, and collagen I. We found that the cells grew into 3D structures when plated on BME and we did not observe cells with a morphology reminiscent of AT1 cells. In contrast, the cells formed a thin 2D monolayer when plated on invasin or collagen I and reached diameters of up to 100 um, similar to AT1 cells in vivo (figure S3F).

### ALVO comparison to scRNA-Seq lung tissue data emphasizes regenerative phenotype of ALVOs

To further characterize the organoid-derived cells and compare these to their tissue counterparts, we performed 10x Genomics scRNA-Seq on three organoid lines (Lu37, Lu38, Lu39) grown in ALVO-EM, AT2-MM, and AT1/2-M (figure 4A). We examined the expression level of a published gene signature for AT2 and AT1 cells for each culture condition^30^. Consistent with their expected phenotypes, AT2-MM cultures showed the highest expression of AT2 marker genes, followed by ALVO-EM, while AT1/2-M showed reduced expression. Conversely, the AT1 signature was enriched in the AT1/2-M condition and reduced in the other two conditions (figure 4B; table S2).

**Figure 4:**
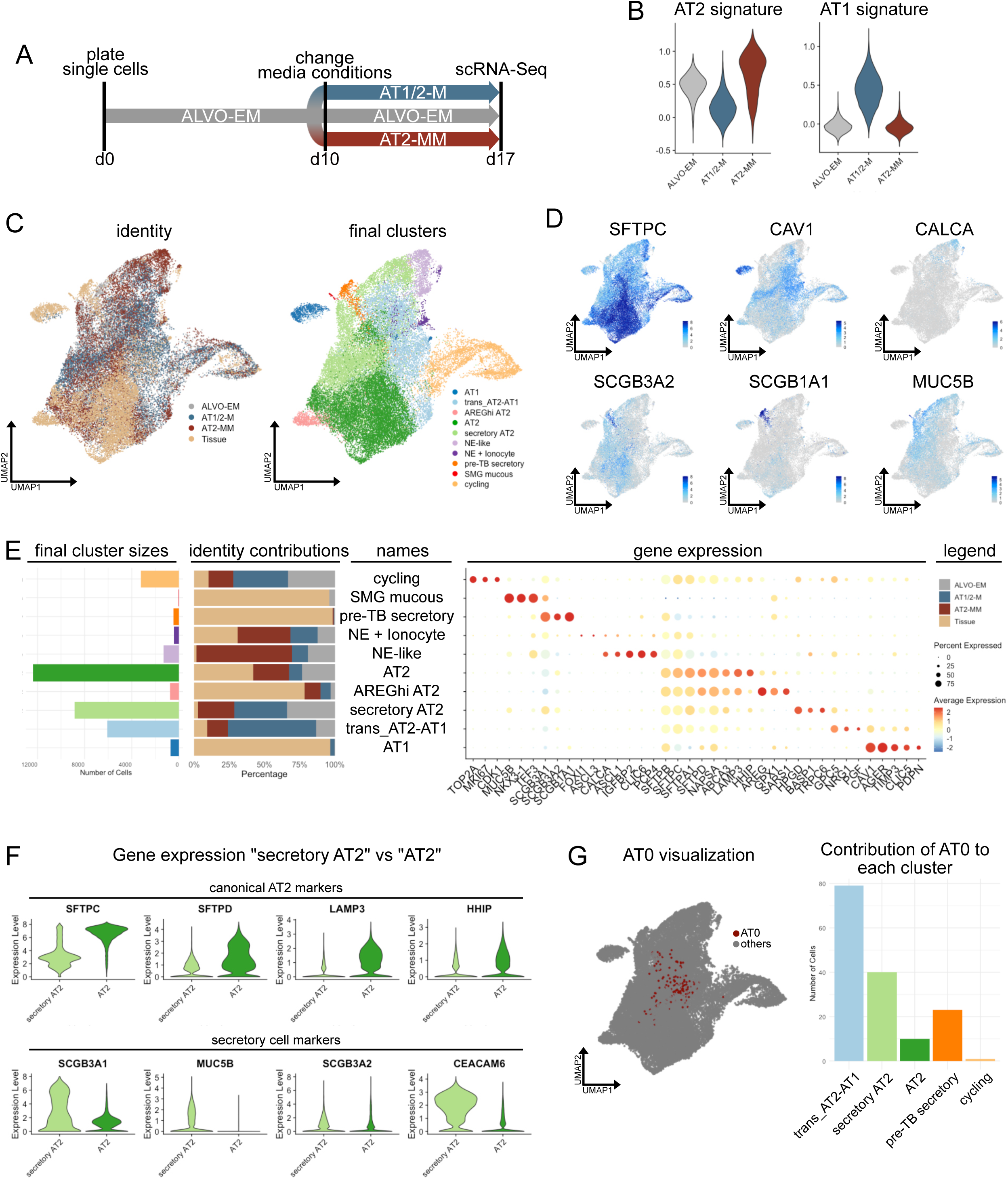
ALVO comparison to scRNA-Seq lung tissue data emphasizes regenerative phenotype of ALVOs. A) Time line of media conditions used for p5 ALVO cultures that were processed for scRNA-Seq. B) Violin plots showing expression levels of published AT2 and AT1 signatures by media condition. C) UMAP showing integrated organoid/tissue subset, colored by organoid media condition and tissue origin (identity, left plot) and seurat clusters (final clusters, right plot). D) UMAP showing expression levels of indicated genes. E) Integrated organoid/tissue subset. Left: Bar plot showing absolute cell contributions to each final cluster. Center: Bar plot showing relative cell contributions of the identities to each cluster. Right: Dot plot showing gene expression of selected genes. F) Violin plots showing gene expression levels of selected genes in “secretory AT2” and “AT2” clusters. G) UMAPs visualizing the location of AT0 cells (from tissue annotations) (left), and bar plot showing the contribution of AT0 cells to final clusters (right) See also figure S4.

To contextualize these findings, we integrated our dataset with the epithelial cell cluster from a previously published lung tissue dataset^33^ (figure S4A). Using the annotations of the tissue data set, we observed that the organoid-derived cells overlapped with alveolar cell populations including “AT0”, “AT1”, “AT2”, and “AT2 proliferating”. Additionally, these cells appeared in close proximity to “Neuroendocrine” (NE), “SMG mucous” (SMG: submucosal gland), and “pre-TB secretory”. Unsupervised clustering identified 31 clusters in total. For subsequent analysis, we selected clusters that included organoid-derived cells (figure S4A).

The resulting subset was re-clustered and annotated based on known marker genes and prior tissue annotations (figure 4C-E; table S3; see methods for more details). All major clusters contained contributions from all three organoid lines (figure S4B). Among these, the “cycling” cluster was highly enriched for proliferation-associated genes (*TOP2A, MKI67, CDK1*) and comprised predominantly organoid-derived cells from all three conditions, underscoring the pro-proliferative environment of organoid cultures relative to homeostatic lung tissue (figure 4E). The “SMG mucous” cluster, containing mostly tissue-derived cells, expressed known SMG goblet cell markers (*MUC5B, NKX3-1, TFF3*). The “NE + Ionocyte” cluster included tissue-derived ionocytes (*FOXI1, ASCL3*) and NE cells (*CALCA, ASCL1*), and also featured contributions from organoids, particularly from the AT2-MM condition (figure 4E and S4C). Interestingly, organoid-derived cells from the AT2-MM condition also formed the majority of an “NE-like” cluster (figure 4E and S4C). Further differential expression analysis of these NE-containing clusters revealed that while ionocyte markers were absent in organoids, NE markers and AT2 markers (*SFTPA2, SFTPC*) were expressed at higher levels (figure S4D). Instead of alveolar markers, tissue-derived cells in this cluster expressed *SCGB1A1*, a club cell marker reported in a subset of NE cells^34^.

The “AT2”, “AREGhi AT2”, and “secretory AT2” clusters exhibited high expression of canonical AT2 markers (*SFTPC, SFTPA1, SFTPD, NAPSA, ABCA3, LAMP3, HHIP*) (figure 4E). The “AT2” cluster, the largest in this subset, contained both tissue- and organoid-derived cells, with most organoid cells derived from ALVO-EM and AT2-MM conditions. Differential expression analysis between tissue- and organoid-derived AT2 cells revealed similar expression levels of canonical AT2 markers (figure S4E; table S3). Notably, genes that differed between these two groups included histone-modifying proteins and ATPases. GO term analysis for biological processes (BP) revealed terms surrounding RNA and protein regulation, possibly reflecting differences of the in vitro versus in vivo environments (figure S4F). Consistent with this interpretation, performing differential gene expression analysis on the entire tissue and organoid cell subset yielded similar findings (figure S4G+H; table S3). The “AREGhi AT2” cluster consisted mostly of tissue-derived cells and likely represents an aberrant AT2 state, as AREG expression has been associated with fibrosis^35^ (figure 4E). “Secretory AT2” cells, compared to the “AT2” cluster, expressed higher levels of the mucin *MUC5B* and the secretoglobin *SCGB3A1*, typically associated with goblet and airway secretory cells, respectively (figure 4E+F). This cluster also showed lower expression of canonical AT2 markers compared to the “AT2” cluster (figure 4F). Moreover, “secretory AT2” cells exhibited higher levels of *SCGB3A2* and *CEACAM6*, a surface marker used for sorting of RAS cells^10^ (figure 4F). The “AT1” cluster, expressing canonical AT1 markers (*CAV1, AGER, TIMP3, CLIC5, PDPN*), contained primarily tissue-derived cells and a small fraction of AT1/2-M organoid cells (figure 4E). Most AT1/2-M organoid cells fell into a “trans_AT2-AT1” cluster, representing a transitional state. This cluster co-expressed both AT1 and AT2 markers, albeit at lower levels than the canonical “AT1” and “AT2” clusters. Because AT0 cells have also been described as an intermediate state, we investigated their location and found that most tissue-derived AT0 cells were present in the “trans_AT2-AT1” cluster, followed by the “secretory AT2” cluster (figure 4G).

In summary, while our organoids contain AT2 cells closely resembling their tissue counterparts, they also contain NE-like, secretory, and transitional alveolar cells. These latter populations are transcriptionally similar to described multipotent progenitors, reflecting the heightened plasticity in organoid cultures compared to homeostatic lung tissue.

### IFN-γ is cytotoxic to AT1-like cells but promotes AT2 growth in a dose- and time-dependent manner

To study alveolar epithelial regeneration in a pro-inflammatory environment as found in COPD, we cultured organoids in ALVO-EM (mostly containing AT2-like cells) or AT1/2-M (containing AT2-like and AT1-transitioning cells) conditions in the presence of cytokines that are upregulated in lungs of COPD patients: IFN-γ, tumor necrosis factor-alpha (TNF-α), IL-6, IL-1α, and IL-1β^36–39^. We omitted the AT2-MM condition in these studies, as the presence of the inflammatory response suppressor dexamethasone could have confounded our results. All tested cytokines led to similar sized or larger organoids in ALVO-EM conditions, while the organoids in AT1/2-M appeared significantly smaller in the presence of IFN-γ but not the other cytokines (figure S5A).

Intrigued by the distinct effect of IFN-γ on the two organoid types, we quantified these changes using a cell viability assay. When IFN-γ was added at the start of organoid culture and maintained for 14 days, cell numbers increased under ALVO-EM conditions but significantly decreased under AT1/2-M conditions (figure 5A). In contrast, IFN-γ added to the organoids on day 11 and incubating for 3 days did not result in cytotoxicity in ALVO-EM conditions, while the cytotoxic effect doubled in AT1/2/M compared to the control (figure 5B). When we examined expression levels of *SFTPA1* (AT2 marker) and *AGER* (AT1 marker) in AT1/2-M organoids, we found that exposure to IFN-γ led to an increase in *SFTPA1* and a decrease in *AGER*, indicating a selective loss of AT1-like cells but not AT2 cells in this condition (figure 5C). To determine if we could rescue the observed cytotoxic effect of IFN-γ in AT1/2-M organoids, we tested the Janus Kinase 1/2 (JAK1/2) inhibitor Ruxolitinib (RX). JAK1/2 acts downstream of the IFN-γ receptor to phosphorylate and activate STAT1, the key transcription factor in IFN-γ signaling. First, we assessed RX tolerance and found that concentrations up to 1 μM were well tolerated, whereas 10 μM was toxic (figure S5B). At 1 μM, RX fully rescued AT1 cells from IFN-γ-induced cytotoxicity and restored expression of *AGER* and *CAV1* to levels comparable to the control (figure 5D, 5E, and S5C). To confirm that this represented on-target effects, we analyzed gene expression levels of the STAT1 target genes *IRF1* and *SOCS1* and found that RX suppressed the expression of these genes in a dose-dependent manner (figure S5D).

**Figure 5:**
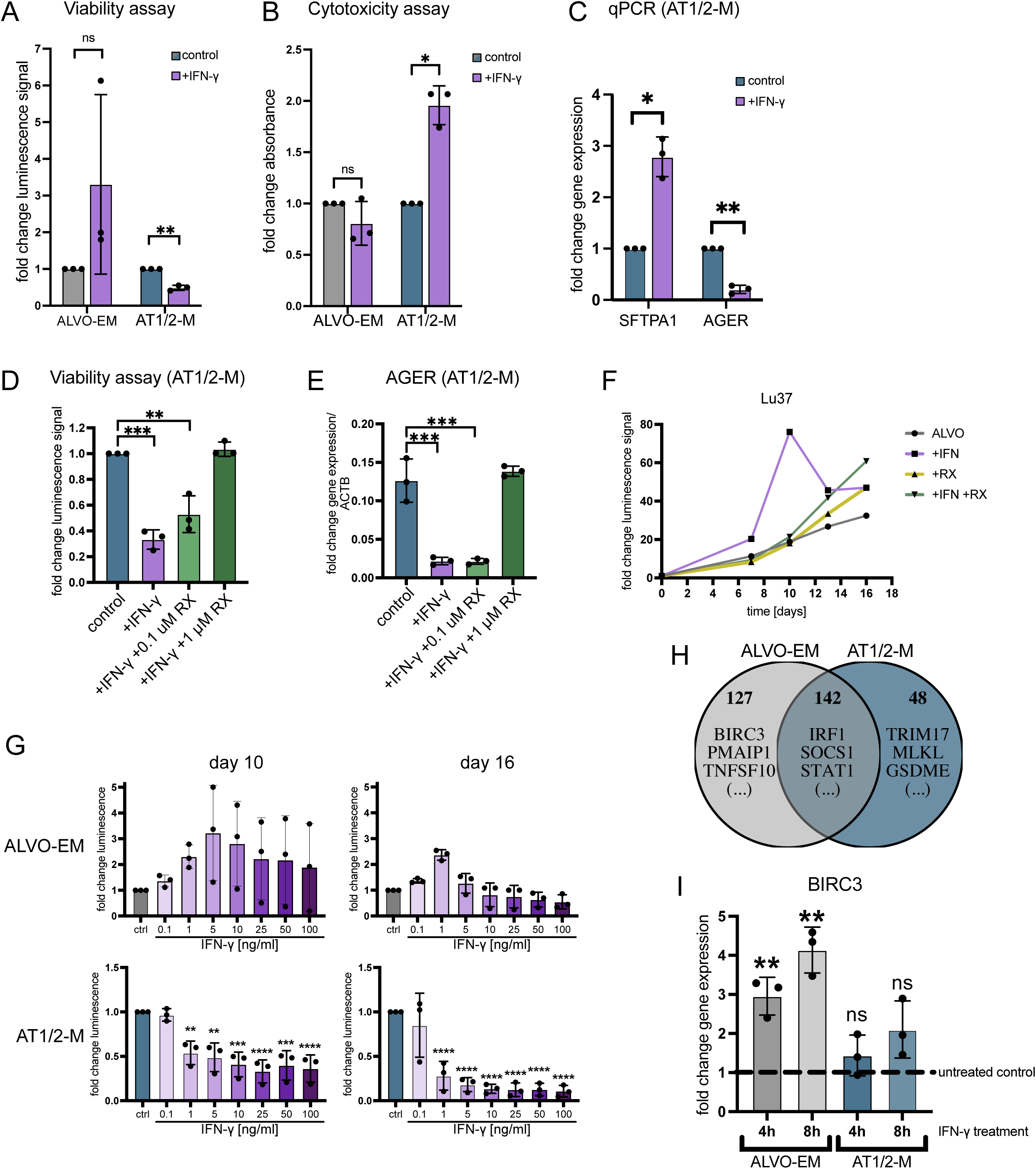
IFN-γ is cytotoxic to AT1-like cells but can promote AT2 growth in a dose- and time-dependent manner. A) Viability assay (cell titer glo) of ALVOs cultured in indicated media conditions in the absence (control) and presence of IFN-γ (10 ng/ml) for 14 days. Data are represented as mean ± SD. B) Cytotoxicity assay of ALVOs cultured in indicated media conditions for 14 days. IFN-γ (10 ng/ml) was added on day 11 for 3 days. Data are represented as mean ± SD. C) qPCR analysis of alveolar markers in ALVOs cultured in AT1/2-M in the absence (control) or presence of IFN-γ (10 ng/ml) for 14 days. Data are represented as mean ± SD. D) Viability assay (cell titer glo) of ALVOs cultured in AT1/2-M in the absence (control) and presence of IFN-γ (10 ng/ml) and RX (0.1 uM and 1 uM) for 14 days. Data are represented as mean ± SD. E) qPCR analysis of AGER in ALVOs cultured in AT1/2-M in the absence (control) and presence of IFN-γ (10 ng/ml) and RX (0.1 uM and 1 uM) for 14 days. Data are represented as mean ± SD. F) Viability assay time course of Lu37 ALVOs cultured in ALVO-EM in the absence and presence of IFN-γ (10 ng/ml) and RX (1 uM). G) Viability assay (cell titer glo) of ALVOs cultured in indicated media conditions for 10 or 16 days in the presence of different IFN-γ concentrations. Data are represented as mean ± SD. H) Venn diagram of genes upregulated in ALVO-EM and AT1/2-M upon treatment of IFN-γ (10 ng/ml) for 4h determined by RNA-Seq. I) qPCR analysis of BIRC3 in ALVO-EM and AT1/2-M upon treatment of IFN-γ (10 ng/ml) for 4h and 8h normalized to their respective untreated controls (dotted line). See also figure S5.

Because the positive effect of IFN-γ on AT2 cell growth showed greater variability among biological replicates than its negative effect on AT1-like cells (figure 5A), we performed a time course in ALVO-EM conditions to understand AT2 growth dynamics in the presence of IFN-γ and RX. Across three biological replicates, organoids exhibited accelerated growth during the first 10 days of IFN-γ exposure, followed by a reduction in cell numbers (figure 5F and S5E). RX treatment prevented both the initial IFN-γ-driven growth spurt and the subsequent decline. The reduction in cell numbers after day 10 was likely due to increased cell death observed in the organoid cultures exposed to IFN-γ (figure S5F).

Our findings indicate that short-term IFN-γ treatment promotes the growth of AT2 cells, whereas prolonged exposure leads to cell death. To determine whether the distinct effects of IFN-γ on AT1-like and AT2 cells is dose-dependent, we treated AT1/2-M and ALVO-EM organoids with IFN-γ concentrations ranging from 0.1 to 100 ng/ml. In AT1/2-M cultures, we observed reduced cell numbers at both early (day 10) and late (day 16) time points starting at concentrations as low as 1 ng/ml, suggesting that even low levels of IFN-γ are toxic to AT1-like cells (figure 5G). In contrast, under ALVO-EM conditions, low IFN-γ concentrations promoted cell growth, while toxicity emerged only at higher doses (figure 5G). Although this pattern was consistent across all three biological replicates, the degree of variation indicates that different donors may exhibit varying sensitivities to IFN-γ. In summary, IFN-γ negatively affects AT1-like cells across a range of concentrations and exposure times, whereas lower concentrations and shorter treatments can enhance the regenerative capacity of AT2 cells.

To better understand the distinct effect of IFN-γ on AT1 and AT2 cells at a molecular level, we treated 14-days-old organoids grown in ALVO-EM or AT1/2-M with IFN-γ and collected RNA at 0.5h, 1h, 4h, and 8h post-treatment. IFN-γ target genes *IRF1*, *SOCS1*, and *STAT1* were upregulated in both media conditions at 4h and 8h of IFN-γ exposure but not at the earlier time points (figure S5G). To further investigate IFN-γ induced changes in gene expression, we performed RNA-Seq on samples collected at the 4h time point. After filtering the cells, we found 142 genes that were upregulated upon IFN-γ treatment in both ALVO-EM and AT1/2-M, including known IFN-γ target genes such as *IRF1*, *SOCS1*, and *STAT1*. In addition, 48 genes were upregulated exclusively in AT1/2-M, including genes associated with cell death such as *TRIM17*, *MLKL*, *GSDME*, and *STING1*^40–43^. 127 genes were uniquely upregulated in ALVO-EM, including genes that modulate cell death, such as *BIRC3*, *PMAIP1*, and *TNFSF10*^44–46^ (figure 5H; table S4). Although *PMAIP1* and *TNFSF10* are generally considered pro-apoptotic, cell death regulation depends on the balance between pro- and anti-apoptotic signals. Notably, only ALVO-EM organoids increased expression of the anti-apoptotic gene BIRC3 at 4h and 8h post-IFN-γ exposure, hinting at a mechanism by which AT2 cells could evade the cytotoxic effects of IFN-γ (figure 5I).

### ALVO-macrophage (MΦ) co-cultures recapitulate the effect of IFN-γ on alveolar epithelial cells

To determine whether the distinct effects of IFN-γ on AT2 and AT1-like cells holds up in a more complex set-up, we developed an ALVO-MΦ co-culturing system that allowed us to test the effect of MΦ-produced cytokines on ALVOs. IFN-γ is produced during viral and bacterial infections by various immune cells, including MΦ, that are themselves stimulated by IFN-γ^47^. MΦs are closely associated with alveolar epithelial cells and are found in greater numbers in COPD patients^48,49^. To model the environment during an immune response to infection, we treated peripheral blood mononuclear cell (PBMC)-derived MΦs with lipopolysaccharides (LPS) and IFN-γ for 24h before setting up a co-culture with ALVOs growing in BME domes in the upper compartment of a transwell (figure 6A). By transferring the transwells into plates containing fresh pre-treated MΦ every 3-4 days, we maintained a consistent cytokine supply for 14 days without having to disrupt the attached MΦs (figure 6B).

**Figure 6:**
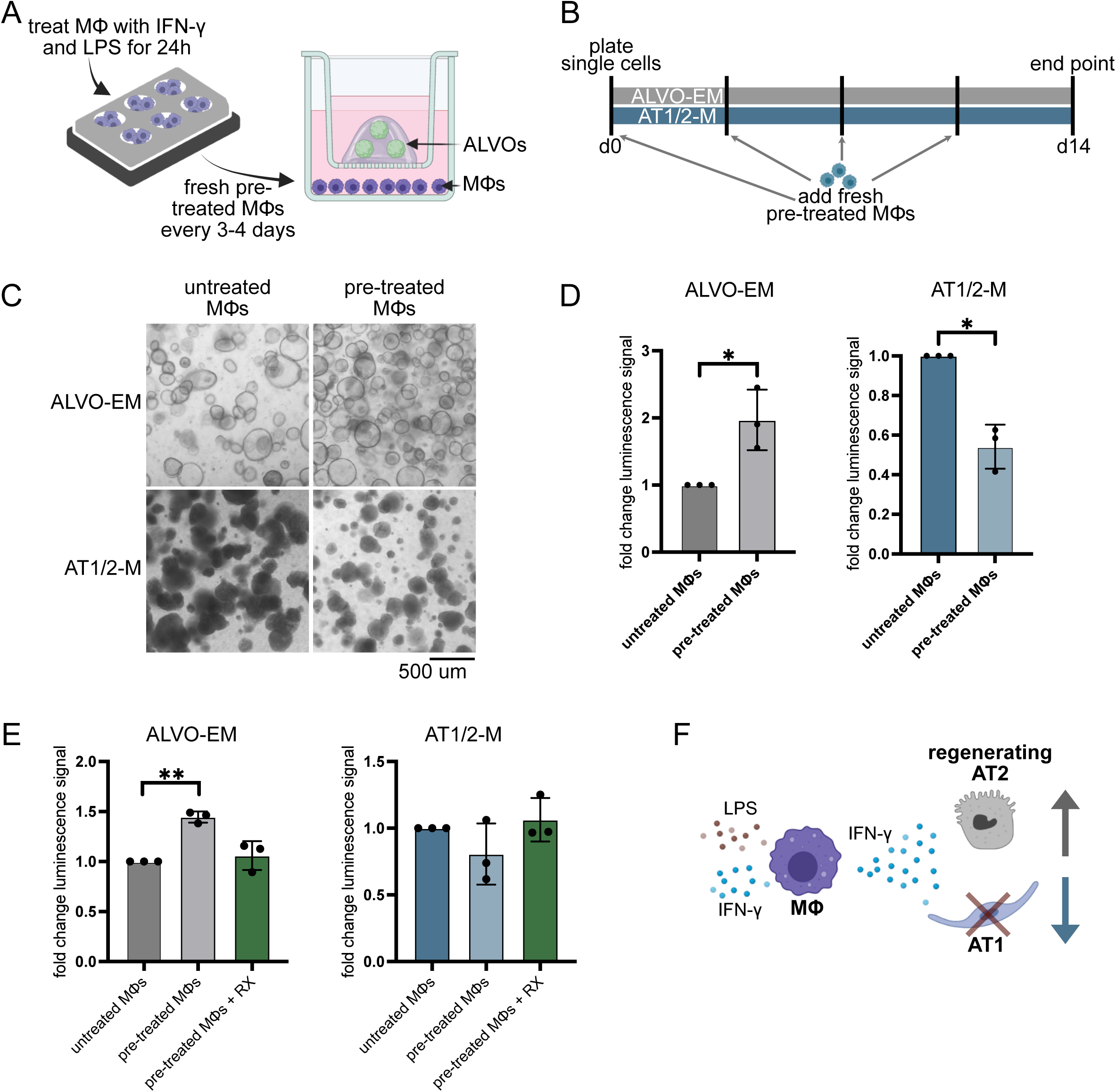
ALVO-MΦ co-cultures recapitulate the effect of IFN-γ on alveolar epithelial cells. A) Schematic of MΦs-ALVO co-culture setup. MΦs are treated with IFN-γ and LPS 24h before they are put into co-culture with ALVOs growing on top of a transwell. Both compartments contain organoid media. B) Time line of MΦs-ALVO co-culture setup. Pre-treated MΦs are added fresh every 3-4 days for a 14 day period. Co-cultures are either exposed to ALVO-EM or AT1/2-M. C) Representative brightfield images of ALVOs in indicated media conditions in the presence of untreated or pre-treated MΦs. D) Viability assay of ALVOs cultured in indicated media conditions in the presence of untreated or pre-treated MΦs. Data are represented as mean ± SD. E) Viability assay of ALVOs cultured in indicated media conditions in the presence of untreated MΦs, pre-treated MΦ, or pre-treated MΦ + RX (1 uM). Data are represented as mean ± SD. F) Summary of macrophage co-culture findings: Macrophages in the presence of LPS and IFN-γ, representing an infection environment, secrete IFN-γ in concentrations sufficient to have a positive effect on regenerating AT2 cells and a negative effect on AT1 cells.

Under these conditions, we observed that MΦ-derived factors induced by LPS and IFN-γ inhibited growth of organoids cultured in AT1/2-M, while promoting the growth of organoids in ALVO-EM, mirroring the direct IFN-γ treatments (figure 6C and 6D). To confirm that these effects were IFN-γ-dependent, we repeated the co-culture in the presence of the IFN-γ signaling inhibitor RX. The addition of RX completely rescued the observed phenotype (figure 6E). In summary, these co-culture experiments show that MΦ exposed to a pro-inflammatory infection-like environment produce cytokines, including IFN-γ, in sufficient concentrations to modulate AT2- and AT1-like cell growth—mediating a growth advantage and growth inhibition, respectively (figure 6F).

## Discussion

There is growing evidence that the notion of AT2 cells as the stem cells of the alveoli that give rise to AT1 cells over-simplifies the alveolar self-renewal process. Lung epithelial cells demonstrate remarkable plasticity, with multiple populations, including airway cells, capable of repopulating the alveolar region following injury^9,10,13,14,50^. To better capture this complexity, we have developed a defined approach to modeling alveolar regeneration with human organoids. The sorting strategy and media conditions facilitate the expansion of cells co-expressing transitioning and secretory markers while maintaining baseline AT2 marker expression. This approach acknowledges the notion that regenerative alveolar progenitor cells are not equivalent to homeostatic AT2 cells. Notably, the co-expression of multiple lineage markers may signify enhanced cellular plasticity and a greater potential to differentiate into diverse lung epithelial cell types.

Under these regeneration-mimicking conditions, the addition of DCI induces AT2 maturation, as evidenced by increased pulmonary surfactant production and the presence of highly ordered tubular myelin in organoid lumens. To our knowledge, this is the first time tubular myelin has been observed in human alveolar in vitro cultures. While dexamethasone is known to accelerate AT2 maturation in fetal lungs, it is mostly used as a systemic anti-inflammatory drug in adults^27,51^. Our findings suggest that dexamethasone may still enhance surfactant production in regenerating adult lungs, raising important considerations for its use as an immunosuppressant in COPD and other lung diseases. Moreover, we observe evidence of an NE-like phenotype in the DCI-containing condition. This finding was unexpected, as no known connection exists between DCI and the induction of a neuroendocrine lineage. Recently, a human fetal alveolar organoid model also found evidence for NE-like cells in their DCI-containing cultures^52^. Further experiments are needed to determine whether rare NE-like cells expand in response to DCI or if DCI can induce NE-like features.

YAP signaling has recently emerged as a key factor of AT1 differentiation. Our results show that YAP activation via LATS inhibition is sufficient to promote an AT1-like phenotype. However, the majority of AT1-like cells have a gene expression profile resembling the AT2-to-AT1 transitional state rather than that of fully differentiated AT1 cells. It is possible that organoid-derived differentiated AT1 cells are underrepresented in the scRNA-Seq dataset due to technical reasons. The typical shape of AT1 cells makes this cell type especially sensitive to dissociation methods necessary to achieve single-cell suspensions. A biological explanation for the underrepresentation is that additional signals or prolonged culture periods might be required for terminal AT1 differentiation. Nevertheless, we posit that the presence of transitional states in our organoids better reflects regenerative conditions and provides a valuable platform for optimizing AT1 differentiation in therapeutic contexts.

MUC5B is expressed in SMGs and the distal airways of human lungs, typically restricted to club cells under homeostatic conditions^53^. However, MUC5B expression in AT2 cells has been observed in disease, particularly in idiopathic pulmonary fibrosis^54,55^. In ALVOs, we identified a subpopulation of cells co-expressing AT2 markers and *MUC5B*, along with elevated *SCGB3A1* and *CEACAM6* expression, markers linked to secretory cells^9,10^. This suggests that MUC5B expression in these cells may represent an adaptive feature of alveolar regeneration, potentially amplified by organoid pro-proliferative and stemness-inducing organoid culture conditions. Whether this co-expression occurs physiologically during alveolar repair or represents a culture artifact requires further investigation.

Our findings on IFN-γ provide insights into the complex interplay between tissue repair and inflammation during acute infections in emphysema. IFN-γ, a key cytokine primarily released by T cells in response to viral and bacterial infections, is elevated in COPD patients compared to healthy controls^39,56^ and further increases during infection-triggered exacerbations^57,58^. The literature on the direct effect of IFN-γ on alveolar epithelial cells is conflicting. While one study found that IFN-γ promoted both AT2 proliferation and differentiation to AT1 cells^59^, another study found that the addition of IFN-γ led to increased cell death in alveolar spheroid cultures^29^. This discrepancy may partly arise from differences in the IFN-γ concentrations used: in vitro studies commonly apply ng/ml concentrations, whereas physiological estimates for IFN-γ levels during acute and chronic lung inflammation fall within the pg/ml range^60,61^. In our study, IFN-γ exhibited a consistently negative impact on AT1-like cells across a variety of concentrations tested, while its effect on regenerating AT2 cells varied depending on dose and exposure duration, with lower concentrations promoting growth of the cells. Notably, we demonstrated that IFN-γ secreted by co-cultured stimulated MΦ was sufficient to replicate these effects, adding physiological relevance to our findings. These results imply that blocking IFN-γ signaling to prevent tissue destruction in emphysema patients may inadvertently impair AT2 regeneration. A deeper understanding of the pathways that render AT1 cells vulnerable and protect AT2 cells in IFN-γ-rich environments could inform the development of therapeutic strategies to enhance alveolar repair while effectively managing inflammation.

In conclusion, our study presents a thoroughly characterized ALVO model for investigating alveolar regeneration under inflammatory conditions. A key strength of this model is its controlled design, where the sole variable between AT2- and AT1-promoting conditions is the addition of LATS-i. This feature minimizes confounding factors and enables precise mechanistic studies. Our platform holds promise for advancing drug screening efforts to identify factors that promote alveolar regeneration and AT1 differentiation, particularly in the context of inflammatory diseases such as COPD.

### Limitations of study

Unlike many other human adult organoid systems, ALVOs are still limited in long-term growth. This makes experiments that involve genetic engineering and require clonal outgrowth challenging, limiting the tools available for precise mechanistic studies. Furthermore, organoids cannot fully recapitulate the complexity of the in vivo lung environment, including mechanical forces, interactions with other cell types, and systemic immune responses.

## Supporting information

Excel file containing AT1 and AT2 gene signatures from Burgess et al. Related to figure 4 and S4.

Excel file containing scRNA-Seq DEGs. Related to figure 4 and S4.

Excel file containing upregulated genes from bulk RNA-Seq. Related to figure 5 and S5.

Excel file containing list of oligos used for gene expression analysis. Related to Materials and Methods

## Acknowledgments

We thank the Singe Cell Genomics core of the Princess Maxima Center (Utrecht, The Netherlands) and the Flow Cytometry Core at the Hubrecht Institute (Utrecht, The Netherlands) for their technical assistance. A.F.M.D. has received funding from the European Respiratory Society (ERS) and the European Union’s H2020 research and innovation programme under the Marie Sklodowska-Curie grant agreement No 847462. The authors are solely responsible for its content; it does not represent the opinion of ERS and the European Commission and ERS and the EU Commission are not responsible for any use that might be made of data appearing therein. This work was also supported by the Netherlands Organ-on-Chip Initiative, an NWO Gravitation project (024.003.001) funded by the Ministry of Education, Culture and Science of the government of the Netherlands (KB, HC), the Oncode Institute (partly financed by the Dutch Cancer Society) (HC) and the Accelerate Lung Regeneration Consortium BREATH (12.0.18.002) of the Lung Foundation Netherlands (LMR, HC). CPC is financially supported by the Gravitation Program “Materials Driven Regeneration”, funded by the Netherlands Organization for Scientific Research (024.003.013).

## Author contributions

Conceptualization: A.F.M.D. Data curation: C.P.C. and G.J.F.S. Formal analysis: A.F.M.D. and C.P.C. Funding acquisition: A.F.M.D., J.H.E, and H.C. Investigation: A.F.M.D., K.B., L.M.R., W.E., W.W., C.L.I., and H.B. Project administration: A.F.M.D., J.H.E., and H.C. Resources: N.S. Supervision: A.F.M.D., P.J.P., and H.C. Validation: A.F.M.D. and L.M.R. Visualization: A.F.M.D. Writing – original draft: A.F.M.D. Writing – review & editing: A.F.M.D., K.B., C.P.C., L.M.R., and H.C.

## Declaration of generative AI and AI-assisted technologies in the writing process

During the preparation of this work, the authors used ChatGPT-4o to spell- and language-check written text. After using this tool, the authors reviewed and edited the content as needed and take full responsibility for the content of the publication.

## Materials

| REAGENT or RESOURCE | SOURCE | IDENTIFIER |
| --- | --- | --- |
| **Antibodies** |  |  |
| anti-NGFR-PE | BioLegend | Cat#345106 |
| anti-EPCAM-APC | BioLegend | Cat#369810 |
| anti-AGER | R&D Systems | Cat#AF1145 |
| anti-SCGB1A1 | R&D Systems | Cat#MAB4218 |
| anti-GPRC5A | abbexa | Cat#abx005719 |
| anti-HT2-280 | Terrace Biotech | Cat#TB-29AHT2-280 |
| anti-KRT5 | Biolegend | Cat#905501 |
| Anti-SFTPC | Santa Cruz | Cat#sc-518029 |
| **Biological samples** |  |  |
| Lung tissue | Diakonessenhuis Utrecht |  |
| PBMCs | Sanquin Amsterdam |  |
| **Chemicals, peptides, and recombinant proteins** |  |  |
| AdDMEM/F12 | Gibco | Cat#12634010 |
| Pen/Strep | Gibco | Cat#15140-122 |
| HEPES | Gibco | Cat#15630-056 |
| L-GlutaMAX | Gibco | Cat#35050-038 |
| Primocin | InvivoGen | Cat#Ant-pm-1 |
| B27 | Gibco | Cat#17504-044 |
| N-Acetylcysteine | Sigma-Aldrich | Cat#A9165 |
| Heparin | Sigma-Aldrich | Cat#H3149 |
| CHIR9902 | Tocris | Cat#4423 |
| SB431542 | Selleckchem | Cat#S1067 |
| BIRB796 | MedChemExpress | Cat#HY-10320 |
| FGF-10 | R&D systems | Cat#345-FG/CF |
| HRG1-b1 | Peptotech | Cat#100-03 |
| Dexamethasone | Sigma-Aldrich | Cat#D4902 |
| cAMP | Sigma-Aldrich | Cat#B7880 |
| LATS-i | MedChemExpress | Cat#HY-138489 |
| EGF | Peptotech | Cat#AF-100-15 |
| Y-27632 | AbMole Bioscience | Cat#M1817 |
| IL-1b | Biolegend | Cat#579402 |
| Collagenase | Sigma-Aldrich | Cat#C9407 |
| DNase | Sigma-Aldrich | Cat#DN25 |
| Red blood cell (RBC) lysis buffer | Roche | Cat#11814389001 |
| Basement membrane extract (BME) | R+D Systems | Cat#3533-005-02 |
| TrypLE | Gibco | Cat#2605010 |
| FBS | Sigma-Aldrich | Cat#F7524 |
| DAPI | Sigma-Aldrich | Cat#10236276001 |
| LysoTracker Green | Fisher Scientific | Cat#11594976 |
| Nile Red | Sigma-Aldrich | Cat#72485 |
| IFN-γ | Peptotech | Cat#300-02 |
| Ruxolitinib (RX) | MedChemExpress | Cat#HY-50856 |
| DAPT | Sigma-Aldrich | Cat#D5942 |
| FGF7 | Peprotech | Cat#100-19 |
| Human serum (HS) | Sigma-Aldrich | Cat#H4522 |
| IL-1A | Peprotech | Cat#200-01A |
| IL-6 | Peprotech | Cat#200-06 |
| IL-33 | Peprotech | Cat#200-33 |
| IL-4 | Peprotech | Cat#200-04 |
| IL-10 | Peprotech | Cat#200-10 |
| IL-13 | Peprotech | Cat#AF-200-13 |
| IL-22 | Peprotech | Cat#200-22 |
| Invasin | Wijnakker et al.<sup>33</sup> |  |
| Collagen I | Advanced Biomatrix | Cat#5074 |
| Dispase | Gibco | Cat#17105041 |
| Phalloidin | Sigma-Aldrich | Cat#65906 |
| Lymphoprep | Stemcell Technologies | Cat#18061 |
| Cellbanker freezing media | Amsbio | Cat#11888 |
| RPMI 1640 medium | Gibco | Cat#61870010 |
| M-CSF | Peprotech | Cat#300-25 |
| *E. coli* LPS | Sigma-Aldrich | Cat#L4391 |
| *P. aeruginosa* LPS | Sigma-Aldrich | Cat#L9143 |
| 1,4-Dithiothreitol | Sigma-Aldrich | Cat#D9779 |
| PneumaCult™ Alveolar Organoid Expansion Media | Stemcell Technologies | Cat#100-0847 |
| PneumaCult™ Alveolar Organoid Differentiation Medium | Stemcell Technologies | Cat#100-0861 |
| DRAQ5 | Biostatus | Cat#DR05500 |
| Critical commercial assays |  |  |
| Cell titer glo (CTG) assay | Promega | Cat#G9683 |
| Cytotoxicity assay | Promega | Cat#G1780 |
| RNA extraction kits | Quiagen | Cat#74106 ,<br>Cat#74536 |
| High-Capacity RNA-to-cDNA kit | Applied Biosystems | Cat#16271891 |
| iQ SYBRGreen supermix | BioRad | Cat#1725006CUST |
| Deposited data |  |  |
| scRNA-Seq data | GEO | GSE287209 |
| Oligonucleotides |  |  |
| List of all oligonucleotides | See table S5 |  |
| Software and algorithms |  |  |
| GraphPad Prism 10 |  |  |
| R (v.4.4.1) |  |  |
| CellRanger (v.7.2.0) | Zheng et al.<sup>64</sup> |  |
| SoupOrCell algorithm (v2.5) | Heaton et al.<sup>65</sup> |  |
| Seurat package (v.5.1.0) | Hao et al.<sup>66</sup> |  |
| DESeq2 package (v.1.44.0) | Love et al.<sup>63</sup> |  |
| Other |  |  |
| 500 µm filter | pluriSelect | Cat#43-50500-03 |
| 100 µm filter | Greiner bio-one | Cat#542000 |
| Suspension culture plates | Greiner bio-one | Cat#657185,<br>Cat#665102,<br>Cat#662102 |
| white-walled 96-well plates | Greiner bio-one | Cat#655098 |
| black-bottom, black-walled 96-well plate | Greiner bio-one | Cat#655076 |
| Transwell inserts | Greiner bio-one | Cat#662641,<br>Cat#665641 |
| Cell scrapers | Greiner | Cat#541070 |

## Methods

### Human samples

Distal human lung tissue samples were obtained as healthy adjacent tissues from tumor resections at the Diakonessen Hospital Utrecht (Utrecht, The Netherlands). Informed consent was acquired from all donors. The study was approved by the ethical committee and was in accordance with the Declaration of Helsinki and according to Dutch law. This study is compliant with all relevant ethical regulations regarding research involving human participants. As patient samples were anonymized, sex, gender, age, race and other information were not recorded and are not available.

### Tissue processing for organoid cultures

Non-tumor lung tissue from lobectomies were kept in saline solution on ice or in washing media (table S1) at 4°C until processing. Tissues were transferred to a sterile dish and minced using scissors and razor blades. The mechanically dissociated tissue was transferred to a 50 ml conical tube and 10 ml prewarmed digestion buffer (washing media + 0.5 mg/ml collagenase (Sigma-Aldrich)) was added per gram of tissue, but not exceeding 25 ml per tube. Tissue was incubated in the digestion buffer on a shaker at 37°C for 25 minutes (min). Following incubation, DNase (50 ug/ml; Sigma-Aldrich) was added and tubes were placed on ice for 5 min. The digested tissue was filtered through a 500 µm filter (pluriSelect) placed on a 50 ml tube, using the flat end of a syringe plunger to push the tissue through the filter. The filter was rinsed with 10 ml of cold washing media, and the process was repeated with a 100 µm filter (Greiner bio-one). Filtered suspensions were centrifuged at 400 g for 5 min, and supernatants were carefully aspirated without disturbing the cell pellet. Cell pellets were resuspended in ∼1 ml of red blood cell (RBC) lysis solution (Roche) per gram of tissue and incubated at room temperature (RT) for 5 min. The lysis reaction was quenched with 15 ml of washing buffer (table S1), the cell suspension was centrifuged at 400 × g for 5 min, and the supernatant was aspirated. If the cell pellet was still red, the RBC lysis step was repeated no more than once. The resulting cell pellet typically has a color ranging from white to dark grey, depending on the amount of pollution present in the tissue. The pellet was resuspended thoroughly in 1 ml of washing buffer (table S1). If large clumps were observed, the suspension was filtered through a 100 µm filter (Greiner bio-one) and spun down again. The number of live cells was determined using Trypan Blue, and cells were resuspended in a mixture of 20% washing buffer (table S1) and 80% BME (R+D Systems) at a concentration of ∼3000 cells/µl. Drops of 20–40 µL were plated on pre-warmed suspension culture plates (Greiner bio-one). After the BME solidified (10-15 min), pre-warmed organoid media supplemented with Y-27632 (10 uM; AbMole Bioscience) and optionally IL-1β (10 ng/ml; BioLegend) was added for the first few days. Media was changed every 2-4 days.

### Making single-cell suspensions and passaging organoids

Organoids were passaged approximately every 14 days. After aspirating media, ∼1 ml TrypLE (Gibco) was added to one well of a 6 well plate (scale volume down for smaller formats) and BME domes were disrupted mechanically by pipetting up and down and organoid suspension was collected into a 15 ml canonical tube and incubated at 37C for 2-4 min. For difficult-to-dissociate organoid cultures (in the presence of LATS-i), the incubation time was increased in 4 min increments, up to 30 min, with periodical pipetting. Organoids were disrupted by pipetting up and down rigorously, without introducing bubbles, using a 1000 µl pipette tip or a Pasteur glass pipette. The dissociation progress was monitored under a brightfield microscope periodically, and the reaction was quenched by adding 10 ml cold washing media (table S1) when the suspension was mostly single cells. The number of live cells was determined using Trypan Blue. Cells were spun down at 400 g for 5 min and media was aspirated. If a BME layer was visible above the cell pellet, the pellet was resuspended in cold washing media and spun again. When no BME was visible above the cell pellet anymore, the cells were resuspended in a mixture of 20% washing buffer (table S1) and 80% BME (R+D Systems) at a concentration of 300-400 cells/µl. Drops of 20–40 µL were plated on pre-warmed suspension culture plates (Greiner bio-one). After the BME solidified (10-15 min), pre-warmed organoid media supplemented with Y-27632 (10 uM; AbMole Bioscience) was added for the first few days. Media was changed every 2-4 days.

### FACS of p0 organoids

Organoids at p0 were made into single-cell suspensions as outlined in “Making single-cell suspensions and passaging organoids**”** and resuspended in FACS buffer [PBS + 2% FBS (Sigma-Aldrich) + 2 mM EDTA] at 5-10 million cells/ul. Cells were divided into several tubes for fluorophore-minus-one (FMO) staining controls. DAPI (Sigma-Aldrich), anti-NGFR-PE (1:100), and anti-EPCAM-APC (1:100) were added and cells were incubated on ice for 15 min. Cells were spun down at 400 g for 5 min, supernatant was discarded, pellet was resuspended in FACS buffer, and cell suspension was strained into FACS tubes. Cells were gated on single, DAPI-, EPCAM+, NGFR-cells and sorted into ALVO-EM (table S1) culturing media using a BD FACSAria Fusion. Cells were plated in a mixture of 20% washing buffer (table S1) and 80% BME (R+D Systems) at 300-400 cells/ul as outlined in “Making single-cell suspensions and passaging organoids”.

### Flow cytometry analysis of LysoTracker and Nile red stained organoid cells

Organoids that were cultured in ALVO-EM (table S1) for 10 days followed by AT2-MM (table S1) for 7 days were made into single-cell suspensions as outlined in “Making single-cell suspensions and passaging organoids” and resuspended in washing media to a concentration of 1 million cells/ml. DAPI (Sigma-Aldrich) and either LysoTracker Green (100 nM; Fisher Scientific) or Nile Red (500 ng/ml; Sigma-Aldrich) were added, and cells were incubated while rotating at 37C for 45 min or 30 min, respectively. After the incubation time, cells were spun down at 400 g for 5 min, supernatant was removed, and the cells were washed with FACS buffer [PBS + 2% FBS (Sigma-Aldrich) + 2 mM EDTA]. After another spinning step, the cell pellet was resuspended in FACS buffer and strained into FACS tubes. Cells were gated on single, DAPI-cells and the median fluorescent signal of lysotracker ([488] 530/30) and Nile Red ([561] 585/15) was measured on a BD LSRFortessa flow cytometer.

### Viability assay: Cell titer glo (CTG)

Single cells were plated at a density of 300-400 cells/µl in a mixture of 20% washing buffer (table S1) and 80% BME (R+D Systems), with a total volume of 5 µl per well in white-walled 96-well plates (Greiner bio-one). The BME was allowed to solidify for 10–15 min at 37C before adding 100 µl of pre-warmed media. The media conditions containing varying concentrations of IFN-γ (Peprotech) and RX (MedChemExpress) are outlined in the figures and were added from day 0 on, with Y-27632 (10 uM; AbMole Bioscience) present for the first few days. For each condition, 3-4 wells were plated as technical replicates. Media was replaced every 2-4 days. On the days of analysis, the manufacturer’s protocol for the CTG assay (Promega) was followed. Briefly, 100 µl of CTG solution was added to the 100 ul media containing wells (1:1 ratio), and the plate was protected from light and shaken for 10 min before incubating for 5 min. A 100 µl sample of the media/CTG mix was then transferred from each well into a black-bottom, black-walled 96-well plate (Greiner bio-one). Luminescence signals were recorded using a standard plate reader. Measurements on day 0 immediately after plating were done to account for differences in plating between organoid lines. Subsequent measurements were normalized to day 0 values, and normalized again to the control group for each organoid line.

### Cytotoxicity assay of IFN-γ-treated organoids

Cells were plated for the cytotoxicity assay the same way as described for the viability (CTG) assay with 3-4 technical replicates per condition. IFN-γ (10 ng/ml; Peprotech) was added in 100 ul fresh media on day 11 of culture, and the cytotoxicity assay (Promega) was performed on day 14 according to the manufacturer’s protocol. Briefly, 50 ul of media was transferred from each well into a new clear bottom 96 well plate, an equal volume of CytoTox reagent was added, and the plate was shaken for 5 min. The mixture was incubated at RT for 30 min. Subsequently, 50 ul stop solution was added, the plate was shaken for 5 min, and absorbance was measured at 490 nm using a standard plate reader.

### Small molecule, growth factors, human serum, and cytokine treatments

Small molecules LATS-i (10 mM; MedChemExpress), DAPT (1 uM; Sigma-Aldrich), and RX (100 nM, 1 uM, 10 uM; MedChemExpress), growth factors EGF (50 ng/ml; Peprotech) and FGF7 (10 ng/ml; Peprotech), and HS (10%; Sigma-Aldrich) were added to organoid culture according to the timelines stated in the figures. Cytokines IL-1A (Peprotech), IL-1B (BioLegend), IL-6 (Peprotech), IL-33 (Peprotech), IL-4 (Peprotech), IL-10 (Peprotech), IL-13 (Peprotech), IL-22 (Peprotech), and IFN-γ (Peprotech), were added to organoid cultures at a concentration of 10 ng/ml unless stated otherwise.

### Transwell cultures

Transwell inserts (Greiner bio-one) were coated with 10 µg/ml Invasin^32^ in PBS or 2% Collagen I (Advanced Biomatrix) in washing media (table S1) at 4C overnight, or with 5% BME (R+D Systems) in washing media (table S1) at 37C for 30 min. Day 14 organoids cultures were processed to single-cell suspensions as outlined in “Making single-cell suspensions and passaging organoids” and 100k-200k cells were plated per transwell. Media was added to the bottom and top of the transwell until cell layer was confluent, usually after one week. At this point, media was removed from the upper compartment to establish air-liquid interface (ALI) conditions. Transwells were fixed after approximately one week of ALI culture.

### Immunohistochemistry for organoid slides, transwells, and whole-mount samples

Dispase (1 mg/ml; Gibco) was added to the media of the submerged organoids and the plate was incubated at 37C for 45-60 min to dissolve the BME. The organoid suspension was carefully collected into a 15 ml conical tube using a 1 ml pipette tip with cut off tip to prevent shearing of the larger organoids. The tube and tip were pre-coated with FBS (Sigma) to prevent sticking of the organoids to the plastic. Cold washing media (table S1) was added to the tube, and the organoids were spun down at 50g for 3 min. The supernatant was removed and the organoids were fixed with 4% paraformaldehyde for 30-60 min at RT. For transwell cultures, the cells were fixed inside of the transwells without the addition of dispase, and the membrane with cells was removed from the transwell for downstream processing.

To prepare slides, the organoids and transwell samples were dehydrated, paraffin embedded, and sectioned. Standard H&E stainings and immunohistochemistry was performed using antibodies against AGER (1:500; R&D Systems), SCGB1A1 (1:250; R&D Systems), GPRC5A (1:1000; Abbexa), HT2-280 (1:500; Terrace Biotech), KRT5 (1:2000; Biolegend), and SFTPC (1:250; Santa Cruz). Phalloidin (1:1000; Sigma-Aldrich) was used to visualize cell boundaries.

Organoids were prepared for whole mount-staining as described in detail elsewhere^62^. HT2-280 (1:250; Terrace Biotech) was added overnight at 4C, and Nile red (500 ng/ml; Sigma-Aldrich) was added for 30 min at RT.

Whole mount-stained organoids and sections were imaged using a Leica Sp8 confocal microscope. Fluorescent images were processed using LAS X software.

### MΦ-organoid co-cultures and viability assay (CTG)

Human PBMCs were obtained from buffy coats (Sanquin Amsterdam) via density gradient using Lymphoprep (Stemcell Technologies) as described by the manufacturer. PBMCs were then frozen in Cellbanker freezing media (Amsbio) at -80C until further use. To obtain MΦs, the PBMCs were seeded in an RPMI 1640 medium (Gibco) supplemented with 10% FBS (Sigma) and 5% PenStrep (Gibco) atop a standard 20 cm plastic tissue culture dish and left to adhere for at least 2h at 4C and 37% CO_2_. The non-adherent cells were then thoroughly washed away and the remaining monocyte-enriched adherent cells were gently detached using cell scrapers (Greiner). For the co-culture experiments, the monocytes were either seeded at a density of 3×10^5^ or 2×10^5^ cells in 12 or 24 well cell culture plates, respectively. To differentiate monocytes into MΦs, the media was supplemented with 40 ng/ml recombinant human M-CSF (Peprotech) for 7 days, with media refresh every 2-3 days. To polarize them towards their pro-inflammatory phenotype, they were stimulated for 24h with a combination of 100 ng/ml LPS [50 ng/ml *E.coli* LPS (Sigma Aldrich) and 50 ng/ml *P. aeruginosa* LPS (Sigma Aldrich)] and 10 ng/ml IFN-γ (Peprotech). On the day of co-culture, the media was aspirated and the MΦs were washed with PBS, before the organoid media and the organoids in transwells were added to the wells.

To grow organoids in transwells, single-cell suspensions of organoids as outlined in “Making single-cell suspensions and passaging organoids” were plated in a mixture of 20% washing buffer (table S1) and 80% BME (R+D Systems) at 300-400 cells/ul into a transwell insert (Greiner bio-one) in a total volume of 30 ul or 20 ul for 12 or 24 well inserts, respectively. After the BME had solidified, the transwells were transferred to the plate containing the pre-treated MΦs, and organoid culture media [ALVO-EM or AT1/2-M (table S1); with or without 1 uM RX (MedChemExpress)] was added on top of the transwell and to the MΦs containing bottom compartment. The organoids in transwells were transferred to a fresh plate containing pre-treated MΦs every 3-4 days and media was refreshed in both compartments at the same time.

On day 14, the transwells with organoids were transferred to a fresh plate, the media was aspirated from the top and replaced with fresh media. The manufacturer’s protocol for the CTG assay (Promega) was followed. Briefly, an equal volume of CTG solution was added to each transwell, and the plate was protected from light and shaken for 10 min before incubating for 5 min and additional mixing by pipetting up and down. For each transwell, 3-4 100 µl samples of the media/CTG mix were transferred into a black-bottom, black-walled 96-well plate (Greiner bio-one) as technical replicates. Luminescence signals were recorded using a standard plate reader. Measurements on day 0 immediately after plating were done to account for differences in plating between organoid lines. Subsequent measurements were normalized to day 0 values, and normalized again to the control group for each organoid line.

### Transmission electron microscopy

Medium was removed from organoid-containing wells, and cold washing media with 10% FBS was added to the well. The BME was carefully disrupted using a 1 ml pipette tip, and the organoid suspension was transferred to Eppendorf tubes. Tubes were incubated on ice for 30 min, followed by centrifugation at 300 × g for 5 min. The supernatant was discarded. For fixation, the organoid pellet was washed once in fixative buffer (1.5 % glutaraldehyde / 0.067 M cacodylate buffered to pH 7.4 / 1 % sucrose) an then incubated in fresh fixative buffer at RT for 3h. The organoids were washed with washing buffer 2 (0.1 M cacodylate (pH 7.4) / 1 % sucrose) and finally resuspended in fresh washing buffer 2 and kept at 4C until further processing. Then, organoids were incubated in 1% osmium tetroxide and 1.5% K4Fe(CN)6 in 0.1 M sodium cacodylate (pH 7.4) for 1 h at 4° C. Next organoids were dehydrated in ethanol (70%, 90%, up to 100%) infiltrated with Epon resin for 2 days, and finally embedded in the same resin and polymerized at 60 °C for 48 h. Ultrathin sections were cut on a Leica Ultracut UCT ultramicrotome (Leica Microsystems) using a diamond knife (Diatome) and mounted on Formvar-coated copper grids. Subsequent staining of the sections with 2% uranyl acetate in 50% ethanol and lead citrate allowed visualization under a Tecnai T12 Electron Microscope equipped with an Eagle 4k × 4k CCD camera (Thermo Fisher). Images were stitched, uploaded, shared and annotated using Omero and PathViewer.

### RNA extraction

Media was aspirated from the organoid cultures, and RNA lysis buffer with 1,4-Dithiothreitol (40 mM; Sigma-Aldrich) was added directly to the organoid-containing BME domes. RNA extraction was performed using the RNA kits (Quiagen) according to the manufacturer’s protocol.

### cDNA preparation and RT-qPCR analysis

cDNA was prepared using the High-Capacity RNA-to-cDNA (Applied Biosystems) kit according to the manufacturer’s protocol. cDNA was diluted 1:10 with Milli-Q water and RT-qPCR was performed using the iQ SYBRGreen supermix (BioRad) and gene specific primers (table S5) in a CFX Connect Real-Time PCR machine (BioRad). The results were analyzed by normalization to the housekeeping gene ACTB (2^−ΔCT^) and, depending on the experiment, to a reference sample (2^−ΔΔCT^).

### Bulk mRNA sequencing

Bulk RNA-Seq was performed on three organoid lines (Lu37, Lu38, Lu39) cultured in ALVO-EM or AT1/2-M (table S1) and treated with IFN-γ (10 ng/ml; Peprotech) for 4h or untreated. RNA was extracted as described above. For the library preparation, a minimum of 100 ng of total RNA was used per condition. The Utrecht Sequencing Facility (USEQ, Utrecht, The Netherlands) carried out the library preparation using the TruSeq Stranded mRNA polyA kit, followed by sequencing on the Illumina NextSeq2000 platform. Reads were mapped to the Human genome (GRCh38) using STAR mapping software (v. 2.7.5c). Read counts were assigned using featureCounts (v2.03.) from the subreads package. Raw count data were imported into R using the read.csv function. The DESeq2^63^ package (v.1.44.0) was used to create a DESeq dataset. The three lines were treated as replicates, and differentially expressed genes were determined for untreated vs IFN-γ treated organoids for each of the two media conditions. Genes with ≤10 reads across all conditions, a p-adjusted value ≥0.05, or a log2 fold change ≤1 were filtered out.

### 10x Genomics scRNA-Seq

Organoids from three donors (Lu37, Lu38, Lu39), cultured in the three media conditions ALVO-EM, AT2-MM, and AT1/2-M, were made into single-cell suspensions as outlined in “Making single-cell suspensions and passaging organoids” and stained with DAPI (Sigma-Aldrich) and DRAQ5 (Biostatus) for 10 min in FACS buffer [PBS + 2% FBS (Sigma) + 2 mM EDTA]. After the incubation time, cells were spun down, resuspended in FACS buffer, and strained into FACS tubes. Live cells (DAPI-/DRAQ5+) were sorted into PBS using a BD FACSAria Fusion. Cells from the same condition but different donors were mixed in equal parts and 25,000 cells per condition were subjected to droplet-based scRNA-seq using the 10x Genomics platform. Libraries were prepared using the 10x Genomics Chromium 3’ Gene Expression solution v3 on a GEMX machine and sequenced on a NovaSeq6000 (Illumina). Reads were mapped to the Human genome (CRCh38) using CellRanger^64^ software (v.7.2.0). Reads were demultiplexed based on genotype using the SoupOrCell algorithm (v2.5)^65^.

### Single nucleotide polymorphism (SNP) analysis and scRNA-Seq genotype matching

To match the sequenced cells to the three donors (Lu37, Lu38, Lu39) we performed SNP analysis. Oligo ligation assay (OLA) SNP typing (Luminex) was performed following manufacturers recommendations. Probes were designed to target population-wide SNPs with an incidence of around 50% covering all major chromosomes and with coverage in whole genome and exome sequencing as well as in RNA sequencing approaches such as polyA-enriched RNA sequencing or total RNA sequencing. Detailed target information and genomic coordinates can be found at: https://github.com/Hubrecht-Clevers/SNP_genotyping_airway/design.txt. DNA was isolated from the three organoid lines (Lu37, Lu38, Lu39) that were used for scRNA sequencing. OLA probes were used to amplify a genomic target before hybridization to MagPlex-TAG microspheres (Luminex). Signal intensities were measured using a Luminex-MagPix (Luminex). Genotype profiles were generated using an in-house generated script https://github.com/Hubrecht-Clevers/SNP_genotyping_airway/SNP_profiles_processing.R. Genotypes from single-cell samples were extracted from SoupOrCell vcf-files generated during genotype demultiplexing. These genotypes were compared with OLA SNP typing results. Sample IDs were assigned based on Euclidian distance between an OLA SNP typing result and SoupOrCell genotype.

### scRNA-Seq data pre-processing

CellRanger and Souporcell output files were processed using the Seurat^66^ package (v.5.1.0) in R (v.4.4.1). Firstly, cells identified as doublets or with an unassigned identity by Souporcell were removed from further analyses. Transcripts found in less than 3 cells were removed. High-quality cells were subsequently obtained by filtering out cells expressing less than 200 or more than 7500 transcripts, and a mitochondrial gene percentage higher than 10%.

### scRNA-Seq data integration with a healthy tissue dataset

To enable identification of cell types in the organoids and comparison of their transcriptome to in vivo lung tissue, we integrated all organoid datasets with the reference tissue scRNA-seq dataset published by Sikkema et al. (2023), which contains curated cell type annotation^33^. Prior to integration, only epithelial cells were subsetted from the object based on the authors’ annotation (ann_level_1). Thereafter, the object was downsampled to contain the same number of cells as the combined organoid datasets (29,064 cells). Integration of the three organoid datasets [ALVO-EM, AT2-MM, and AT1/2-M) (table S1)] with the downsampled tissue dataset was performed using Reciprocal PCA (RPCA) with the default parameters. After computing PCA dimensions, Uniform Manifold Approximation and Projections (UMAPs) were rendered using dims=1:35.

### scRNA-Seq clustering, subsetting and differential expression analysis

Using Seurat’s FindClusters function with a resolution of 0.8, we obtained 31 cell clusters. Given the available information on the cell identifies assigned by Sikkema et al. (2023), tissue-derived clusters could be easily distinguished. For a deeper characterization of organoid-derived cells, we subsequently subsetted all clusters containing organoid cells (i.e., clusters 0, 1, 3, 4, 5, 7, 8, 9, 11, 16, 17, 21, 23, 24, 26, and 30) (figure S4A). Cluster 28 was exclusively tissue-derived and constituted an impure population of AT2 cells combined with alveolar macrophages (MARCO+), thus it was excluded. Subsetting and re-scaling of the data yielded 20 clusters. Clusters 0, 1, 2, and 3 contained the highest numbers of tissue-derived cells annotated as “AT2”. Therefore, these were merged and labelled as “AT2”. Since cells in clusters 4, 5, and 16 expressed both AT2 (e.g., *SFTPC*) and AT1 markers (e.g., *CAV1*), they were annotated as “trans_AT2-AT1”. Clusters 6, 7, 8, and 9 showed the highest expression of secretory markers such as MUC5B and HPGD, hence why they were assigned the identity “secretory AT2”. Clusters 10, 12 and 15 were annotated as “cycling” cells. Cluster 11 was annotated as “NE-like” given the high expression of common neuroendocrine markers such as CALCA and ASCL1. Cluster 13 was defined as “AREGhigh AT2” cells, and cluster 14 was annotated as “AT1” cells. Finally, clusters 17, 18, and 19 were identified based on their original tissue annotation as “pre-TB secretory”, “NE+Ionocyte”, and “SMG mucous” cells, respectively.Differential gene expression analyses between clusters and samples were performed with FindAllMarkers, using min.pct=0.25, and logfc.threshold=0.25.

### Statistical analysis

Data are presented as mean ± standard deviation, with individual data points shown to represent biological replicates (organoid lines from different donors). Statistical analyses were performed using GraphPad Prism 10. For comparisons between two groups, an unpaired t-test was used, or a paired t-test if the data were normalized to the control group. For comparisons involving multiple groups against a control group, or among multiple groups, a two-way ANOVA followed by Dunnett’s post-hoc test was employed. When data were normalized to the control group, a matched two-way ANOVA with Dunnett’s post-hoc test was performed on log-transformed values to account for baseline differences. Annotation for *p* values or adjusted *p* values if a post-hoc test was used are: \**P(adj)* < 0.05, \*\**P(adj)* < 0.01, \*\*\**P(adj*) < 0.001, \*\*\*\**P(adj)* < 0.0001. The lack of an indicated *p* value or the letters “ns” mean that the data point was not significant.

---

**Figure S1:**
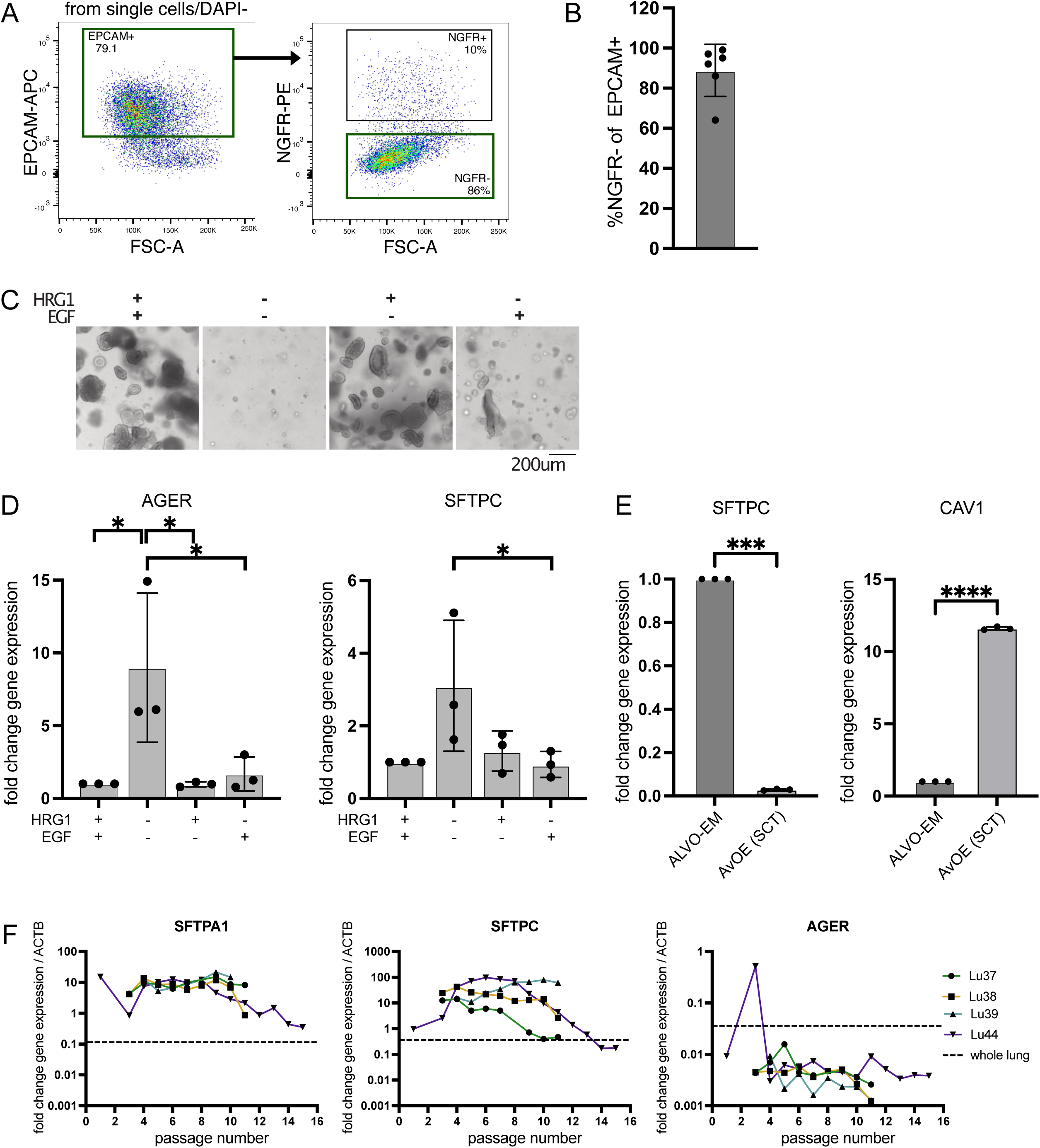
Optimized ALVO conditions allow for long-term expansion in defined media. Related to figure 1. A) FACS plot showing sorting strategy of p0 organoids, including positive selection (indicated by green rectangles) for EPCAM+ cells and NGFR-cells. B) Percentage of NGFR-cells within the EPCAM+ fraction of p0 sorted organoid lines. Data are represented as mean ± SD. C) Brightfield images of p7 ALVO organoids cultured for 14 days with indicated factors. D) qPCR analysis of AT1 (*AGER*) and AT2 markers (*SFTPC*) in ALVOs cultured with indicated factors for 14 days. Data are represented as mean ± SD. E) qPCR analysis of AT1 (*AGER*) and AT2 markers (*SFTPC*) in ALVOs cultured in indicated media conditions for 14 days. Data are represented as mean ± SD. F) qPCR time course of AT2 (*SFTPA1* and *SFTPC*) and AT1 (*AGER*) markers in day 14 old ALVOs at indicated passages with whole lung lysate as reference.

**Figure S2:**
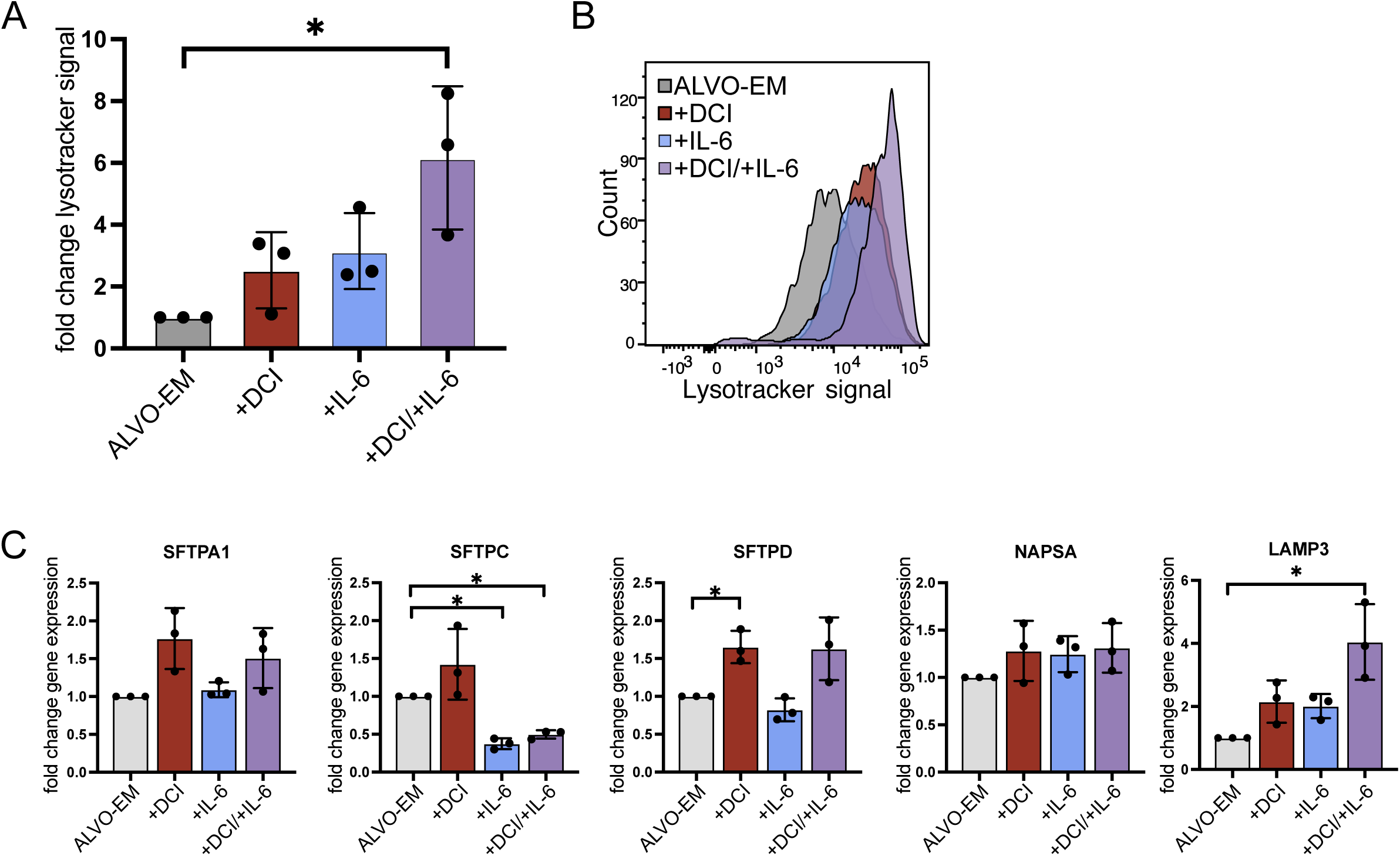
Generating fully mature AT2 cells that secrete tubular myelin-containing surfactant. Related to figure 2. A) Flow cytometry analysis of lysotracker signal in ALVOs cultured in indicated conditions. Data are represented as mean ± SD. B) Representative flow cytometry histogram showing lyostracker signal distribution in indicated conditions. C) qPCR analysis of AT2 markers in ALVOs cultured in indicated conditions as outlined in 2A. Data are represented as mean ± SD.

**Figure S3:**
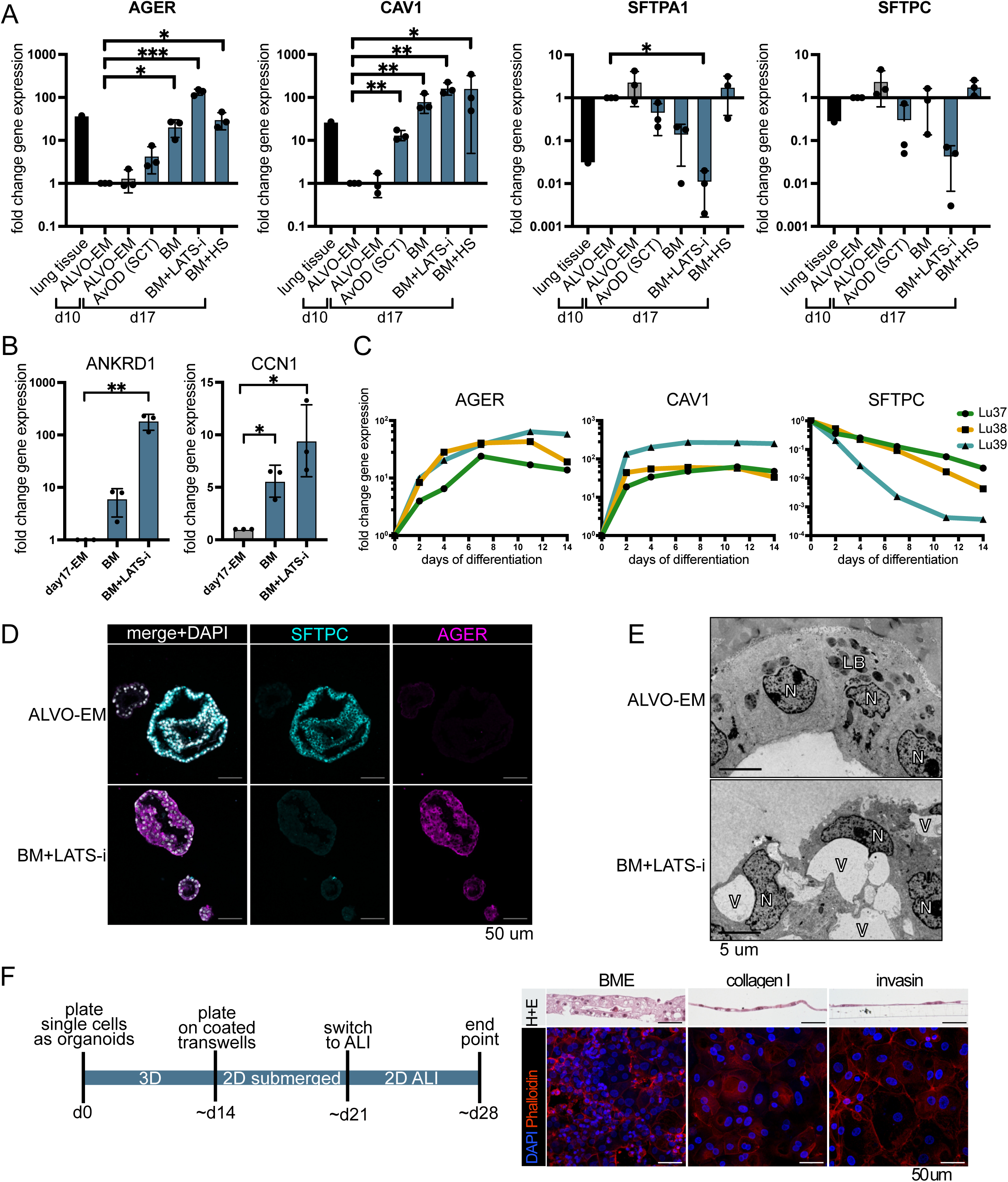
Removal of proliferation and stemness factors in combination with LATS-i drives AT1 phenotype. Related to figure 3. A) qPCR analysis of AT1 (*AGER* and *CAV1*) and AT2 markers (*SFTPA1* and *SFTPC*) in ALVOs cultured in indicated conditions as outlined in 3A with whole lung tissue as reference. Data are represented as mean ± SD. B) qPCR analysis of YAP target genes in ALVOs cultured in indicated conditions as outlined in 3A. Data are represented as mean ± SD. C) qPCR time course of AT1 (*AGER* and *CAV1*) and AT2 markers (*SFTPC*) in the three indicated ALVO lines after switching to BM+LATS-i media following a 10 day expansion phase in ALVO-EM. D) IF images of ALVOs cultured in indicated conditions as outlined in 3A. DAPI = nuclei; SFTPC = AT2 marker; AGER = AT1 marker. E) Electron microscopy image of ALVOs cultured in ALVO-EM or BM + LATS-i as outlined in 3A. N=nucleus; V=vacuole; LB=lamellar bodies. F) Cells were first cultured in 3D, then 2D submerged, then ALI transwells cultures in AT1/2-M as outlined in time line (left). Side-view brightfield (H+E staining) and top-down IF images (right). DAPI = nuclei; Phalloidin = F-actin.

**Figure S4:**
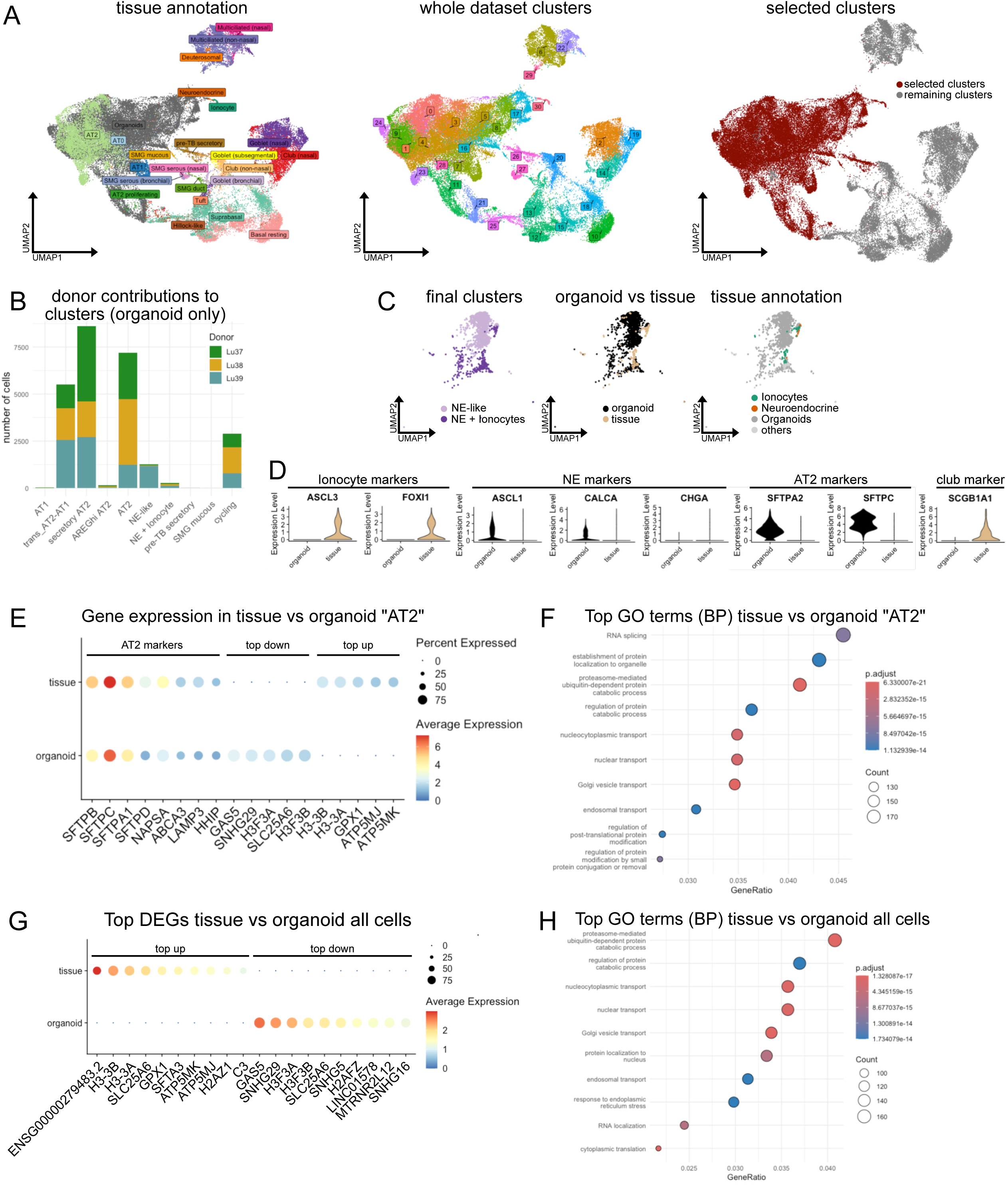
ALVO comparison to scRNA-Seq lung tissue data emphasizes regenerative phenotype of ALVOs. Related to figure 4. A) UMAPs showing integrated organoid/tissue data, colored by tissue annotations and organoids (left), seurat clusters (center), and selected clusters for subsetting (right). B) Bar plot showing the contributions of each organoid line (donor) to the organoid fraction of each final cluster in the organoid/tissue subset. C) UMAP subset of the “NE-like” and “NE + Ionocyte” clusters of the organoid/tissue subset, colored by final cluster (left), organoid vs tissue origin (center), and tissue annotation (right). D) Violin plots showing gene expression levels of selected genes in “NE-like” and “NE + Ionocyte” clusters. E) Dotplot showing AT2 markers and top down- and up-regulated genes comparing organoid and tissue origin within the “AT2” cluster of the organoid/tissue subset. F) Top 10 GO terms (biological processes) of differentially expressed genes between tissue and organoid cells from the “AT2” cluster G) Dotplot showing top up- and down-regulated genes comparing organoid and tissue origin within the whole organoid/tissue subset. H) Top 10 GO terms (biological processes) of differentially expressed genes between tissue and organoid cells from whole organoid/tissue subset.

**Figure S5:**
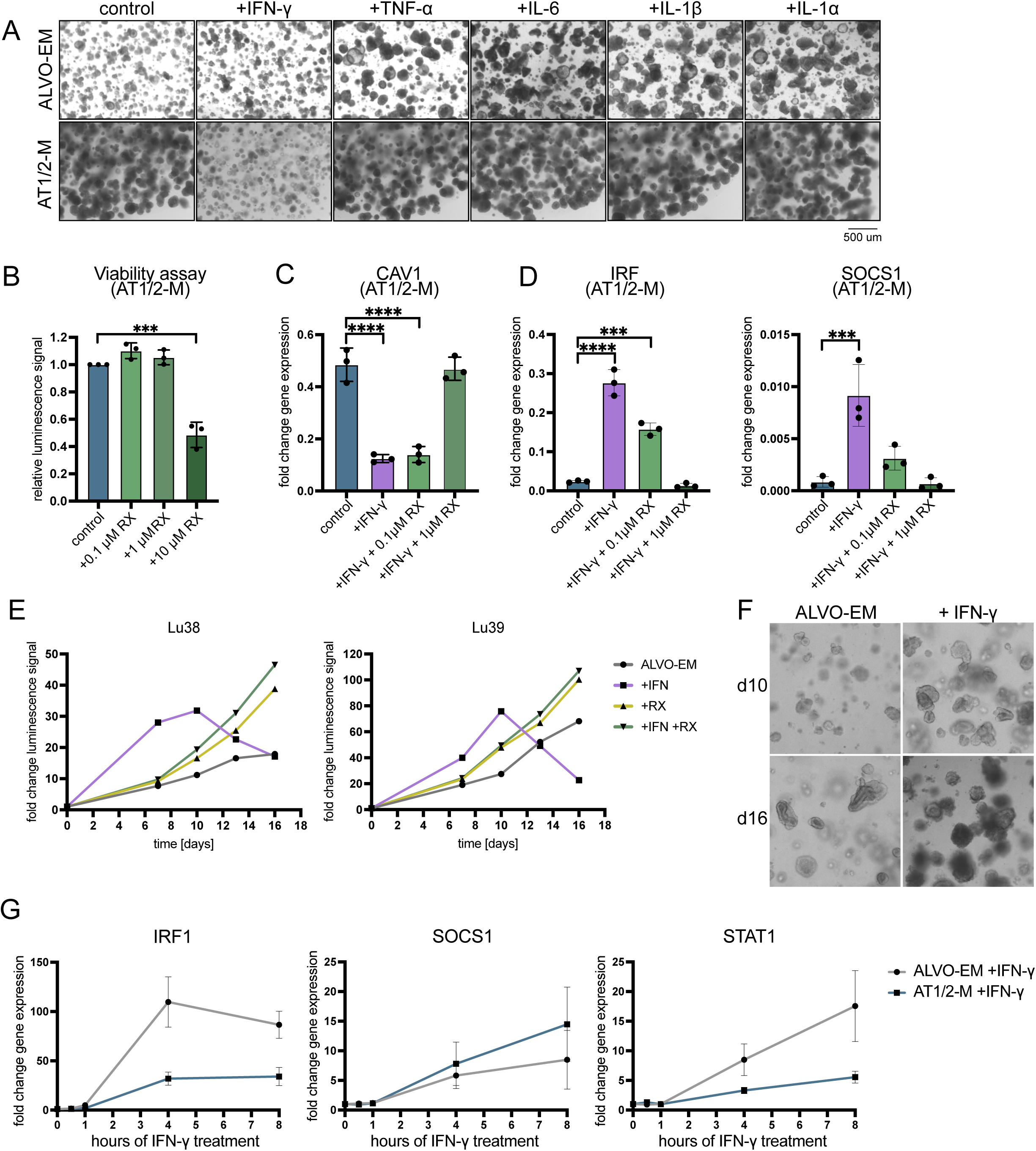
IFN-γ is cytotoxic to AT1-like cells but promotes AT2 growth in a dose- and time-dependent manner. Related to figure 5. A) Brightfield images of ALVOs grown in indicated media conditions in the absence (control) or presence of indicated cytokines for 14 days. B) Viability assay (cell titer glo) of ALVOs cultured in AT1/2-M in the absence (control) and presence of indicated concentrations of RX for 14 days. Data are represented as mean ± SD. C) qPCR analysis of CAV1 in ALVOs cultured in AT1/2-M in the absence (control) or presence of IFN-γ (10 ng/ml) and RX (0.1 uM and 1 uM) for 14 days. Data are represented as mean ± SD. D) qPCR analysis of ALVOs cultured in AT1/2-M in the absence (control) or presence of IFN-γ (10 ng/ml) and RX (0.1 uM and 1 uM) for 14 days. Data are represented as mean ± SD. E) Viability assay time course of Lu38 and Lu39 ALVOs cultured in ALVO-EM in the absence and presence of IFN-γ (10 ng/ml) and RX (1 uM). F) Brightfield images of ALVOs cultured in ALVO-EM in the absence and presence of IFN-γ (10 ng/ml) at indicated time points. G) qPCR time course of IFN-γ target genes in ALVOs cultured in indicated media conditions in the presence of IFN-γ (10 ng/ml) normalized to their respective untreated controls.

**Table S1.** Media compositions. Related to figures 1-6 and S1-5, and STAR methods.

| Component | Concentration | Washing media | BM | SFFF* media (p0) | ALVO-EM | AT2-MM | AT1/2-M |
| --- | --- | --- | --- | --- | --- | --- | --- |
| AdDMEM/F12 | base | + | + | + | + | + | + |
| Pen/Strep | 1x | + | + | + | + | + | + |
| HEPES | 10 mM | + | + | + | + | + | + |
| L-GlutaMAX | 1x | + | + | + | + | + | + |
| Primocin | 100 ug/ml | - | + | + | + | + | + |
| B27 | 1x | - | + | + | + | + | + |
| N-Acetylcysteine | 1.25 mM | - | + | + | + | + | + |
| Heparin | 5 ug/ml | - | + | + | + | + | + |
| CHIR9902 | 3 uM | - | - | + | + | + | + |
| SB431542 | 10 uM | - | - | + | + | + | + |
| BIRB796 | 1 uM | - | - | + | + | + | + |
| FGF-10 | 10 ng/ml | - | - | + | + | + | + |
| HRG1-b1 | 5 nM | - | - | - | + | + | + |
| Dexamethasone | 50 nM | - | - | - | - | + | - |
| cAMP | 100 uM | - | - | - | - | + | - |
| IBMX | 100 uM | - | - | - | - | + | - |
| LATS-i | 10 uM | - | - | - | - | - | + |
| EGF | 50 ng/ml | - | - | + | - | - | - |
| Y-27632 | 10 uM | - | - | day 0-3 | day 0-3 | - | day 0-3 |
| IL-1b | 10 ng/ml | - | - | day 0-3 (p0) | - | - | - |

| absent | present | present on day X |
| --- | --- | --- |
| - | + | day X |

\*from Katsura et al., 2022

